# Harmonizing labeling and analytical strategies to obtain protein turnover rates in intact adult animals

**DOI:** 10.1101/2021.12.13.472439

**Authors:** Dean E Hammond, Deborah M Simpson, Catarina Franco, Marina Wright Muelas, John Waters, R W Ludwig, Mark C Prescott, Jane L Hurst, Robert J Beynon, Edward Lau

**Author notes:** Correspondence: Dean Hammond Edward Lau Robert Beynon. Equal contributions (Dean E Hammond, Edward Lau).

## Abstract

Changes in the abundance of individual proteins in the proteome can be elicited by modulation of protein synthesis (the rate of input of newly synthesized proteins into the protein pool) or degradation (the rate of removal of protein molecules from the pool). A full understanding of proteome changes therefore requires a definition of the roles of these two processes in proteostasis, collectively known as protein turnover. Because protein turnover occurs even in the absence of overt changes in pool abundance, turnover measurements necessitate monitoring the flux of stable isotope labeled precursors through the protein pool such as labeled amino acids or metabolic precursors such as ammonium chloride or heavy water. In cells in culture, the ability to manipulate precursor pools by rapid medium changes is simple, but for more complex systems such as intact animals, the approach becomes more convoluted. Individual methods bring specific complications, and the suitability of different methods has not been comprehensively explored. In this study we compare the turnover rates of proteins across four mouse tissues, obtained from the same inbred mouse strain maintained under identical husbandry conditions, measured using either [^13^C_6_]lysine or [^2^H_2_]O as the labeling precursor. We show that for long-lived proteins, the two approaches yield essentially identical measures of the first order rate constant for degradation. For short-lived proteins, there is a need to compensate for the slower equilibration of lysine through the precursor pools. We evaluate different approaches to provide that compensation. We conclude that both labels are suitable, but careful determination of precursor enrichment kinetics in amino acid labeling is critical and has a considerable influence on the numerical values of the derived protein turnover rates.

## Introduction

Changes in the proteome can be achieved by adjustment of the input into a protein pool (synthesis) or removal of a protein from the pool (degradation), the two processes constituting protein turnover. The simplest model of proteostasis, which is undoubtedly an oversimplification, has three parameters (pool size, synthesis and degradation), linked by zero order synthesis (the rate of synthesis is insensitive to the pool size) and first order degradation (a proportion of the protein pool is degraded per unit time). At steady state, the unchanging pool size is given by the balance between the opposing fluxes of synthesis (molecules/time) and removal (protein pool multiplied by the fractional rate of degradation; thus, also with the dimensions of molecules/time). An adequate description of proteostasis requires that we can measure at least two of these parameters, from which the third can be calculated. Because protein turnover can occur in the absence of any change in pool size, it is evident that turnover parameters must be measured by the flux of a tracer through the protein pool. Most commonly, this is achieved in cells in culture with radiolabeled (e.g. [^35^S]methionine) or stable isotope labeled (e.g. [^13^C_6_]lysine) protein precursors (‘dynamic SILAC’ ^1, 2^). The ability to exchange culture media quickly *in vitro* means that precursor pools can be rapidly manipulated and thus, a transition from labeled to unlabeled media, or *vice versa*, can be made very rapid, relative to protein turnover rates, which minimizes the effects of precursor pool equilibration ^3^.

It is now clear that when compared with cells in culture, protein turnover in animal tissues occurs in completely different temporal regimes, with turnover rates spanning several orders of magnitude. Moreover, different tissues have distinct average turnover rates (for example, liver has a higher turnover rate than skeletal muscle ^4–6^) and larger animals have much lower average rates of protein turnover ^6^. This is in part due to the different energetics constraints between free living animals and cultured cells. The latter grow exponentially in excess nutrients and through cell division can also remove ‘old’ proteins *via* passive dilution, reducing the need for energetically costly proteostatic degradation. This casts doubt on the applicability of cell culture study to understanding turnover in organismal physiology, growth, and aging, and strongly calls for direct measurements of turnover in animal systems.

Unlike cells in culture, in animal systems, the rapid exchange of precursor pools is not always feasible or practical. Isotopically labeled precursors can be administered enterally or parenterally but in both circumstances there is a delay in equilibration of the labeled precursor with the tissue pools, such that in the early phases of labeling, high turnover proteins are sampling a precursor pool that has yet to reach equilibrium. Early studies used radiolabeled amino acid precursors, and although scintillation counting permitted the measurement of very low levels of radiolabel incorporation, this approach was only suitable for total protein pools or measuring purified proteins ^7, 8^. The need to understand proteostasis on a proteome-wide scale has increased the need to measure protein turnover for multiple proteins in the same system, and requires the deployment of stable isotopes. Stable isotope labeling, in combination with proteomics, can yield turnover rates for individual members of the proteome. An additional complication in animal tissues is that turnover rates can be low, and it is difficult to measure very low levels of stable isotope in proteomics-focused mass spectrometry. Thus, labeling duration must be sufficient to lead to discernible incorporation of the label. Stable isotope administration is largely oral, through diet or drinking water and inevitably, this route of administration introduces a delay in equilibration of the precursor with whole-body metabolic pools. This delay can introduce systematic underestimates of rates of turnover, simply illustrated (**Figure 1C**) by modeling of a two-compartment model ^9^.

**Figure 1.**
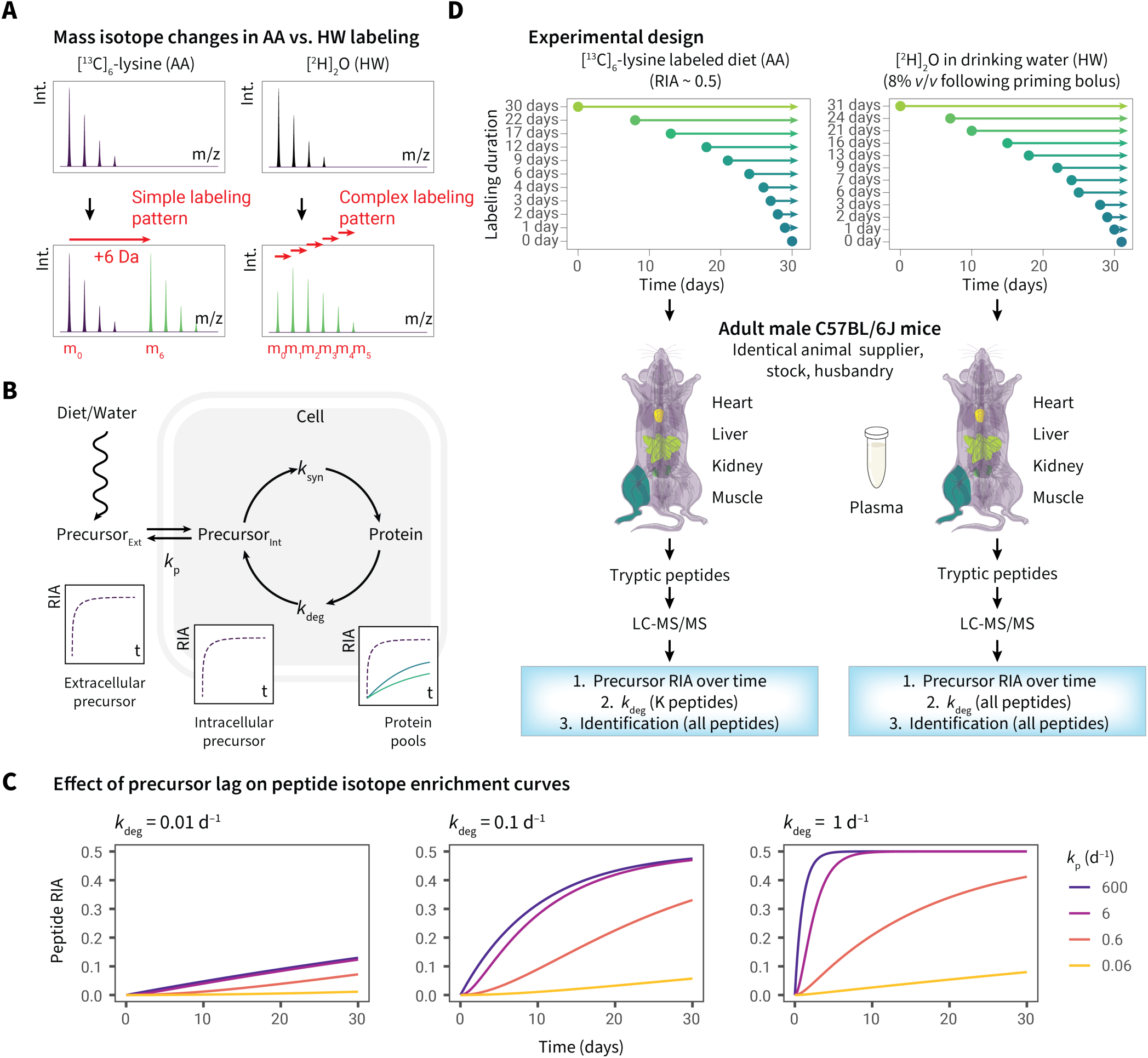
Comparison of labeling strategies for turnover studies in intact adult animals. **A** Mass spectrum features in AA labeling (left) which creates new peptide isotope clusters and elemental HW labeling (right) which shifts the endogenous isotopomer rightward in the mass spectrum. **B.** Schematic of precursor introduction and pool enrichment showing the availability of intracellular precursors for protein synthesis. **C.** The effect on protein labeling of a delay in precursor equilibration. The curves model the effect of a delay in precursor equilibration on labeling of protein pools, for three proteins with degradation rate constants of 0.01 d^−1^, 0.1 d^−1^ and 1 d^−1^ (half lives of 69 d, 6.9 d and 0.69 d, respectively). Four precursor equilibration rates are modelled, with the purple line representing such a high rate (600 d^−1^) as to be equivalent to near-instantaneous equilibration through the body, giving no perceptible delay. As the delay becomes more prolonged, the protein labeling becomes commensurately slower, leading to an underestimate of the true degradation rate constant. **D.** Experimental design. Groups of mice, identical in strain (C57BL/6J), age, sex (male), supplier and husbandry were each labeled with either [^13^C_6_]lysine or [^2^H_2_]O for 30 days, sampling tissues throughout the labeling period. Subsequently, tissues were recovered and tryptic digests were prepared from tissue homogenates to gain protein identity and to assess the degree of isotopic incorporation into proteins.

For animal studies, two approaches are most used, both based on exposure of subjects to stable isotope precursors followed by measurement of the rate of isotope incorporation into individual proteins. First, a labeled essential amino acid can be provided in the diet, either with a relative isotope abundance (RIA) of 1, which requires a fully synthetic diet ^10–12^, or at a lower RIA by supplementation of a standard laboratory diet with pure labeled amino acid ^4, 5, 13, 14^. Typically, the labeled amino acid incorporates multiple heavy atom centers, such that labeled peptides yield *m/z* values that are well resolved from the natural isotope distribution of the unlabeled amino acid (**Figure 1A**, left). Alternatively, animals can be provided with metabolically simple precursors, such as [^2^H]_2_O or [^15^N]H_4_Cl, that deliver a single labeled atom center to some or all amino acids ^15–18^. In this instance, the labeling trajectory leads to a gradual shift of the isotopic profile with considerable overlap between the unlabeled isotopomer profile and the labeled profile (**Figure 1A**, right).

Thus, the incorporation of labels into proteins (and therefore, into peptides derived from those proteins; an essential element in the proteomics workflow) is very different with the two labeling protocols. For amino acid labeling (“AA”) strategies, the incorporation of one or more instances of the labeled amino acid creates ‘heavy’ peptides that are offset by the number of heavy atom centres in the amino acid (such as [^13^C_6_]lysine or [^2^H_7_]valine ^5^). By contrast, for example, the deuterium atoms in heavy water (“HW”) labeling strategies are incorporated stably into specific amino acids, leading to complex labeling patterns wherein labeled peptides have mass shifts from 1 to many Da higher ^17–22^ (**Figure 1A**).

A second difference between the AA and HW strategies pertains to the equilibration of the label with the amino acid pool that is the immediate precursor of protein synthesis. Dietary amino acids need to cross the intestinal mucosal barrier, pass through the hepatic system and are eventually transported to peripheral tissues through the blood (**Figure 1B**). By contrast, water crosses all membranes and is rapidly equilibrated across all tissues ^18^. If equilibration of the label with the precursor is considerably faster than the rate of turnover of the protein pool, then it can be assumed that the precursor enrichment is constant over the labeling period (**Figure 1C**). Under this circumstance, a simple monoexponential function will define the transition from unlabeled to labeled protein. In this regard, HW labeling should equilibrate rapidly, which can be aided by an initial bolus injection of pure [^2^H]_2_O. However, if the precursor pool equilibrates at rates similar to the fastest turnover proteins, then a more complex model is appropriate ^5^. It follows that the AA strategy could be compromised by a delay in pool equilibration, and this would be particularly evident in proteins that were substantially synthesized during the equilibration phase, specifically, high turnover proteins. Because of this unavoidable lag, there have been a number of different solutions to address slow precursor equilibration with amino acids ^9, 11, 12, 23, 24^.

To explore the differences between the AA and HW strategies and to attempt to harmonize the two approaches, we compared the turnover profiles of multiple proteins, derived from four tissues, in mice that were otherwise identical in genotype, source, age, sex and husbandry (**Figure 1D**). This study allowed us to compare the two labeling approaches with a precision not previously realized. Here we present the outcomes of these experiments and show that whilst each approach yields quantitatively comparable results for slow turnover proteins, they are increasingly discrepant for high turnover proteins in a simple exponential kinetics model. In particular, a HW methodology seems to consistently yield turnover rate constants that are higher than those obtained by an AA strategy. When two-compartment models are used to correct for the delay in equilibration of the labeled precursor(s), the rate constants converge more closely.

## Materials and Methods

Both labeling studies were conducted with laboratory mice derived from the same supplier and maintained under identical conditions. Fully grown adult male C57BL/6JOlaHsd mice (obtained from Harlan UK Ltd, Shardlow, UK at 6-13 weeks of age) were previously group housed and used in non-invasive behavioral studies. At the start of this experiment, males aged 15-16 months old were housed individually in 48 x 15 x 15 polypropylene cages (NKP Cages Ltd, Coalville, UK). Each cage contained substrate (Corn Cob Absorb 10/14; IPS Ltd, London, UK), paper wool nest material and environmental enrichment (hanging baskets, plastic tubes). Food (LabDiet 5002 Certified Rodent Diet, Purina Mills, St. Louise, USA) and water were provided *ad libitum*. The mice were maintained on a reversed photo-period (light 12h; dark 12h; lights on at 20:00 hrs) and at 19–21 °C. Animal use and care was in accordance with EU directive 2010/63/EU and UK Home Office code of practice for the housing and care of animals bred, supplied and used for scientific purposes. Heavy water labeling was carried out under UK Home Office licence under the Animals in Scientific Procedures Act 1986 (PPL 40/3492). The University of Liverpool Animal Welfare Committee approved the work.

### Labeling with [^13^C_6_]lysine

This study used eleven mice. Standard laboratory diet (LabDiet 5002) was supplemented with pure, crystalline [^13^C_6_]lysine (Cambridge Isotope Laboratories) to bring the relative isotope abundance (RIA) to 0.5. The dietary pellets were dissociated with water containing the dissolved [^13^C_6_]lysine to form a thick paste and mixed extensively. Once homogeneous, the paste was then extruded into strips 1 cm across and dried in a commercial foodstuff drying oven at 40 °C. The mice had access to the labeled diet for varying amounts of time with randomized assignment: 0, 1, 2, 3, 4, 6, 9, 12, 17, 22 or 30 days. The day that the animals were introduced to the labeled diet was staggered for all endpoints so that dissections took place on the same day. All mice were humanely killed on day 30 and the animals were dissected to recover liver, kidney, heart and pooled hindlimb skeletal muscle from each animal. All tissues, and the carcasses, were frozen at –80 °C prior to analysis.

### Labeling with [^2^H_2_]O

For the heavy water labeling protocol, all animals (13) were provided free access to LabDiet 5002. At the start of the experiment, mice were injected with two successive 0.5 mL injections, 4 hours apart, of 0.15 M sodium chloride dissolved in deuterated water. Thereafter, mice were given free access to 8% (v/v) [^2^H_2_]O for the duration of the experiment. After 0, 1, 2, 3, 6, 7, 9, 13, 16, 21, 24 and 31 days, mice were killed and dissected exactly as described for the [^13^C_6_]lysine labeling experiment and tissues were stored at –80 °C prior to analysis. Additionally, plasma samples were obtained by *post-mortem* cardiac puncture.

### Preparation of samples for proteomics

Small portions (typically 50–100 mg wet weight) from the frozen organs from both studies were further cut into small pieces to facilitate homogenization in 1 mL of lysis buffer (7 M urea, 2 M thiourea, 2 % [*w*/*v*] CHAPS, 5 mM DTT) using a Precellys lysis kit (Stretton Scientific Ltd., Stretton, UK). Total protein extracted was quantified using a Bradford assay. Protein (200 µg, AA; 100 µg, HW) was reduced, alkylated and digested with trypsin using a modified version of the filter-aided sample preparation (FASP) approach ^25^. The labeling protocols were designed so that all labeling time points for a single tissue (11 samples, AA; 12 samples HW) were prepared and analysed concurrently.

Non-targeted MS1-DDA analyses were conducted on a Q-Exactive HF quadrupole-Orbitrap mass spectrometer coupled to a Dionex Ultimate 3000 RSLC nano-liquid chromatograph (Hemel Hempstead, UK). One µg of peptides from each time-point were loaded onto a trapping column (Acclaim PepMap 100 C18, 75 µm x 2 cm, 3 µm packing material, 100 Å) using a loading buffer of 0.1 % (v/v) TFA, 2 % (v/v) acetonitrile in water for 7 min at a flow rate of 12 µL min^-1^. The trapping column was in-line to an analytical column (EASY-Spray PepMap RSLC C18, 75 µm x 50 cm, 2 µm packing material, 100 Å) and peptides eluted using a linear gradient of 96.2 % A (0.1 % [v/v] formic acid): 3.8 % B (0.1 % [v/v] formic acid in water:acetonitrile [80:20] [v/v]) to 50 % A:50 % B over 90 min at a flow rate of 300 nL min^−1^, followed by washing at 1% A:99 % B for 8 min and then re-equilibration of the column to starting conditions. The column was maintained at 40 °C, and the eluent was introduced directly into the integrated nano-electrospray ionisation source operating in positive ion mode. The mass spectrometer was operated in data-dependent acquisition (DDA) mode with survey scans between m/z 350–2000 acquired at a mass resolution of 60,000 (FWHM) at m/z 200. The maximum injection time was 100 ms, and the automatic gain control was set to 3e6. The 16 most intense precursor ions with charge states of 2+ to 5+ were selected for MS/MS with an isolation window of 1.2 m/z units. The maximum injection time was 45 ms, and the automatic gain control was set to 1e5. Fragmentation of the peptides was by higher-energy collisional dissociation using a stepped normalized collision energy of 28–30%. Dynamic exclusion of m/z values to prevent repeated fragmentation of the same peptide was used with an exclusion time of 20 sec.

### Experimental measurement of precursor enrichment

Plasma samples (50 µL) were treated with 2 µL of 10 M sodium hydroxide (BDH, Poole, U.K.) and 1 µL of acetone (Fisher, Loughborough, U.K.). After mixing, the samples were left overnight at 20 °C to allow the exchange of deuterium from water to acetone to occur. To produce a calibration curve, 50 µL mixtures of between 0 and 10 % (*v*/*v*) deuterium oxide (Cambridge Isotope Laboratories, Andover, MA, USA) in HPLC grade water (VWR International, Fontenay-sous-Bois, France) were also treated and extracted. The acetone was then extracted from the samples using 200 µL of chloroform (VWR, Poole, UK) for 15 s. Aliquots of the extracts were then analysed by gas chromatography – mass spectrometry (GC-MS) on a Waters GCT Premier gas chromatograph-mass spectrometer (Waters, Wilmslow, UK). The chromatography column employed was a 30 m long, 0.25 mm internal diameter, 0.25 µm film thickness DB-17MS (Agilent J&W, Santa Clara, CA, USA). The carrier gas was helium (BOC, Guildford, UK) at 1 ml min^-1^. The injector was operated in the splitless mode at 220 °C and the injection volume was 1 µL. The oven temperature programme was 60 °C to 100 °C at 20 °C min^-1^ with a 1 min hold, then from 100 °C to 220 °C at 50 °C min^-1^. The mass spectrometer was operated in the positive ion electron ionisation mode with source temperature 200 °C, electron energy 70 eV and trap current 200 µA. Mass spectra were recorded in low sensitivity mode between 40 to 100 *m/z* with a scan time of 0.1 s. The spectral intensities of ions at *m/z* 58 and *m/z* 59 were measured using the MassLynx software supplied with the instrument. Comparison of the ratio of *m/z* 59 to *m/z* 58 for the biological samples against the curve generated from the calibration samples allowed the enrichment of deuterium oxide (HW) to be measured.

### Direct analysis of tissue lysine pools

Tissue homogenates (100 µL) were added to 350 µL methanol (LC-MS grade), cooled to –80 °C and maintained on dry ice during the addition. The mixture was vortexed vigorously and centrifuged at 13,300 rpm for 15 min at 4 °C to sediment proteins. Aliquots were subsequently dried in a vacuum centrifuge and stored at –80 °C until LC-MS/MS analysis. Prior to analysis, samples were resuspended in 52 µL water (LC-MS grade), centrifuged at 13,300 rpm for 15 min at 4 °C to remove any particulates and transferred to glass sample vials. Untargeted HPLC-MS/MS data acquisition was as published ^26–28^. Full-scan MS and data dependent MS/MS data were acquired using a ThermoFisher Scientific Vanquish HPLC system coupled to a ThermoFisher Scientific Q-Exactive mass spectrometer (ThermoFisher Scientific, UK) operating in positive ionisation mode as described ^29^. Raw instrument data (.raw files) were exported to Compound Discoverer 3.1 for deconvolution, alignment and annotation, as described ^29^. The peak areas of [^12^C_6_]lysine and [^13^C_6_]lysine were retrieved and used to calculate the relative isotope abundance.

### Post processing of protein labeling data

A summary of the workflow for data processing is given in **Figure 2A**. Thermo .raw mass spectrum files were converted to the mzML format using ThermoRawFileParser v.1.2.0 ^30^. The centroid MS2 spectra were searched against the Uniprot ^31^ *Mus musculus* reviewed database (retrieved 2021-Apr-27) using Comet v.2020_01rev3 ^32^ with contaminant proteins and decoys appended using Philosopher v.3.4.13 ^33^. Search settings include: 20 ppm peptide mass tolerance, 0.02 fragment bin tolerance, 0/1/2/3 isotope error, trypsin specificity with 1 enzyme terminus (semi-tryptic) and 2 allowed missed cleavages, and +15.9949 methionine variable modifications. Additionally, AA labeling experiments allowed +6.0201 lysine variable modifications. The Comet search results were post-processed and filtered using Percolator v.3.0.5 ^34^ standalone distribution with the -Y, -i 20, and -P DECOY_ arguments. Peptides identified at the 1% FDR (Percolator q-value) threshold were used for downstream analysis.

**Figure 2.**
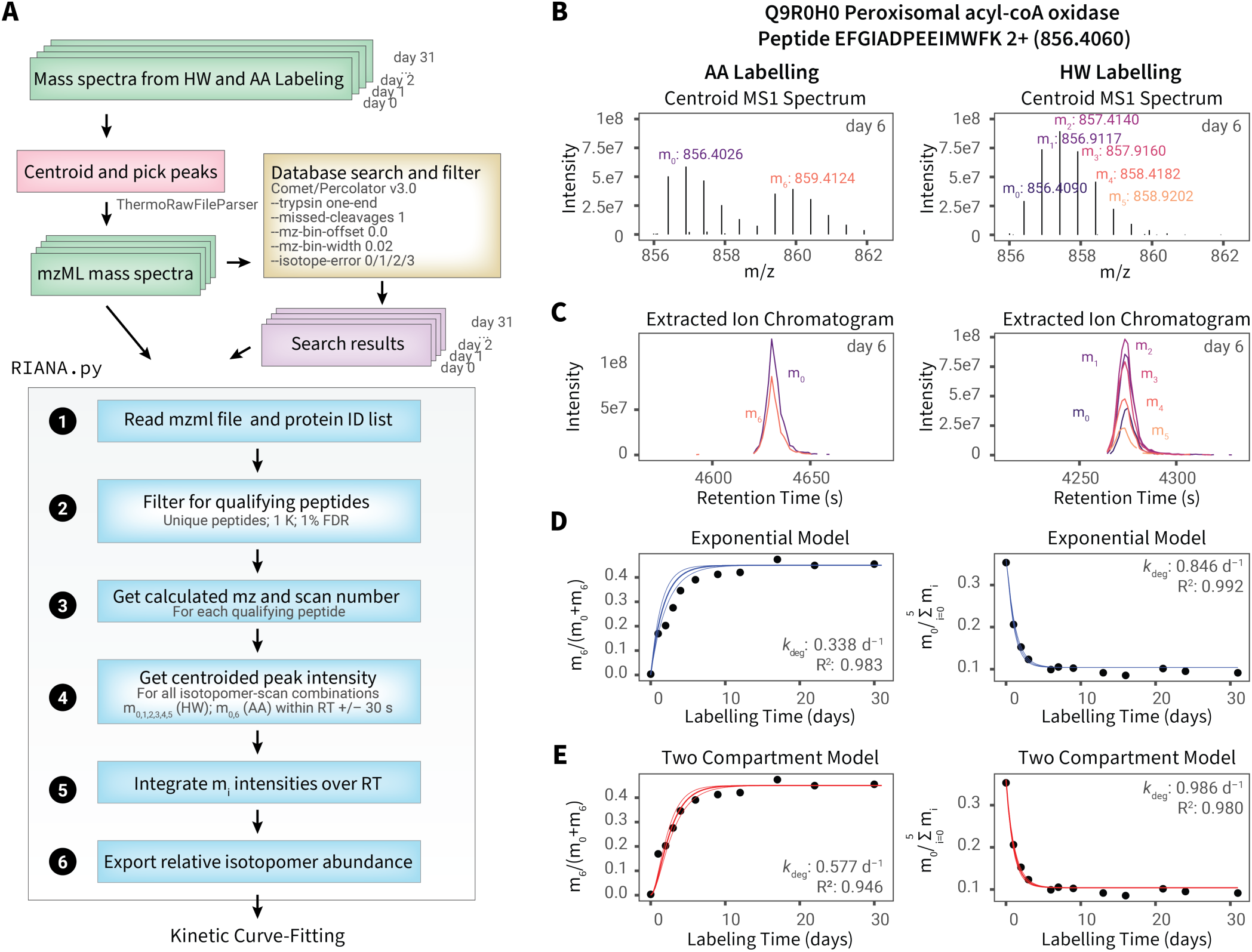
Data analysis workflows. **A.** Schematic for the data analysis workflow for HW and AA labeling data. **B.** The MS1 spectrum for peptide EFGIADPEEIMWFK 2+ of peroxisomal acyl-CoA oxidase (Q9R0H0) for AA labeling (left) and HW labeling (right) data. For AA labeling, the intensity values for the major isotope of light (m_0_) and heavy (m_6_) versions of a peptide are measured for each MS1 scan within a specified retention time window (+/- 30 sec); for HW labeling, the intensities for each successive isotopomer within the isotopomer envelope (m_0_ to m_5_) are measured. **C.** The intensity over time values within the retention time windows are then integrated as shown in the extracted ion chromatograms for AA and HW labeling here. **D.** The data from each labeling time point are processed in the manner described above, resulting in a peptide relative isotope abundance value for each peptide at each time point, which for AA labeling is defined as m_6_/(m_0_+m_6_) and for HW labeling defined as m_0_/(m_0_+m_1_+m_2_+m_3_+m_4_+m_5_). The data time series is then fitted to a simple exponential kinetics model using a quasi-Newton method to optimize for the protein turnover rate constant *k* that results in the least square error value. **E.** Same as panel D, but the time-series data fitted to a two-compartment model to adjust for slow label enrichment in the animal body. The two-compartment model fits the AA data better than the exponential model and leads to a higher estimated *k*_deg_ but has a less pronounced effect on HW labeling due to fast label equilibration.

### Peak integration

To integrate the HW and AA labeling data, we wrote an in-house Python script, Riana v.0.6.4. Riana was written in Python (version >= 3.6) and accepts as input the path to the tab-delimited files generated from Percolator for each organ, labeling method, and experimental time point; and the path to the corresponding directory containing the mzML files to be integrated (**Figure 2A**). Riana uses the pymzml package (≥2.0.0) ^35^ to open mzML files and gathers the intensity values of the centroided peaks of all MS1 spectra for each isotopomer for each qualifying peptide within a retention time range and 25 ppm mass precision (**Figure 2B**). It then integrates and returns the areas-under-curve of the isotopomer chromatograms using the trapezoid method in scipy v.1.6.3^36^. Additional arguments in Riana specify the *n*th isotopomer to be integrated (--iso), which was set to 0,1,2,3,4,5,6,12 for both HW and AA data; the retention time window to integrate across (--rt), which was set to 0.5 min (**Figure 2C**).

### Kinetic models

The collated (RIA, t) series for each peptide charge combination for each fraction in each time point in each organ in each labeling method are then used for kinetic curve-fitting with either a simple exponential (**Figure 2D**) or a two-compartment (**Figure 2E**) model. Kinetic curve-fitting was performed using a custom R script written in R (v.4.1.1) running on platform x86_64-apple-darwin17.0 (64-bit). Optimization for the protein turnover rate constant *k*_deg_ was performed using the optim() function in the stats package of base R using the Broyden-Fletcher-Goldfarb-Shanno (BFGS) quasi-Newton method and a starting value of *k*_deg_ = 0.29. Fitting for the rate constant for precursors to reach plateau (*k*_p_) using the single exponential or Fornasiero double exponential model was optionally also performed using the nls() function with the default Gauss-Newton algorithm to retrieve the log likelihood and calculate the Akaike Information Criterion (prediction error, thus allowing model selection). Time series (*t*, *A_t_*) data were fitted to two models (one-compartment simple exponential vs. two-compartment) to find the best estimate of protein turnover rate (*k*_deg_) that minimizes sums of squares of error. The one-compartment exponential model used is given by:

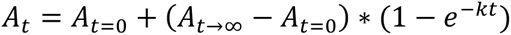

where *A_t_* is the estimated time-dependent relative isotopomer abundance (RIA) of interest for a peptide under a label enrichment level of *p*, at a measured time point *t*, which for HW labeling data is defined as:

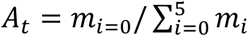

where *m_i_* is the chromatographic area under curve of the *i^th^* isotopomer of the peptide integrated by Riana. *A*_0_ is the initial pre-labeling RIA which for HW is the isotope abundance based on natural isotope distribution calculated from Berglund and Weiser ^37^ and *A_t_*_→∞_ is the asymptotic relative abundance, which for HW was defined by the number of accessible labeling sites at each amino acid based on tritiated water data in Commerford et al. ^38^, the sequence of the peptide, and the deuterium enrichment level as previously described ^15, 17^. For AA labeling, *A_t_* is defined as *m*_6_/(*m*_0_ + *m*_6_) and *A*_0_ is 0% before enrichment which *A_t_*_→∞_ depends on the RIA of heavy amino acid in the feed and can be determined as previously stated. The fitting error of the one-compartment exponential model is given by:

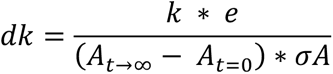

Where *σA* is the residual error of RIA after fitting.

The two-compartment model used is given by:

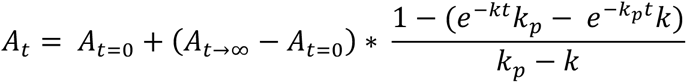

Where *k_p_* is the first-order label accumulation rate constant. The fitting error of the two-compartment model is given by:

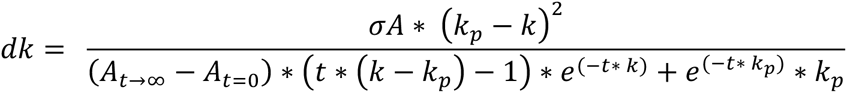

The two-compartment, three-exponent model for peptides and precursor kinetics is as described in Fornasiero *et al.*^11^ For precursor fitting, the model was scaled to 50% heavy lysine.

### Additional data analysis

Data analysis and visualizations were performed in R (v.4.1.1) unless otherwise specified. Robust correlation is performed using biweight midcorrelation implemented in the WGCNA ^39^ package (v.1.70-3). Data visualizations were generated with the aid of the ggplot2 ^40^, gganatogram ^41^, ggpubr ^42^, and plotly ^43^ packages in R. Kernel density estimations were performed using gaussian_kde in scipy (v.1.6.3) ^36^ in Python 3.8. Peptide isotopomer integration output and R code for kinetic curve-fitting have been uploaded to a runnable container at CodeOcean (https://codeocean.com/capsule/3856272/tree/v1).

### Data and code availability

Raw mass spectrometry data have been deposited to ProteomeXchange at PXD029639. The source code and instructions for Riana v.0.6.4 can be accessed at http://github.com/ed-lau/riana.

## Results

For both labeling protocols (AA *vs.* HW), peptides followed the expected trajectory of gradual incorporation of label into peptides. For the AA protocol, the expected increase in intensity of [^13^C_6_]lysine terminated peptides led to clear separation between the unlabeled and labeled components of the peptide pool for all discernible isotopomers (m_0_, m_0_+6; m_1_, m_1_+6 etc.). There was no evidence for partial loss of single labeled atom centres from the amino acid; the isotopomer profiles for labeled or unlabeled peptides are identical. For the HW strategy, the mass shift for the peptide was more subtle, evidenced as a gradual shift from the monoisotopic m_0_ pool and increased intensity of the m_1_, m_2_…m_n_ isotopomer intensities, reflecting gradual incorporation of deuterium into the peptides (**Figure 1A**).

AA-labeled peptides conform to the kinetic model only when the peptide contains one lysine; for fair comparison we initially filtered peptides in the HW experiment identically. With this filter, the HW and AA labeling experiments yielded similar numbers of quantifiable peptides over the entire labeling curve (**Figure 3A–B**). As we have observed previously ^4^, liver and kidney give the highest number of peptides, with heart intermediate and the lowest, skeletal muscle, providing about half as many peptides. We attribute this to the more pronounced dynamic range in protein expression in the two muscle tissues, such that the column loading and analysis are dominated by those proteins that are strongly expressed. There is a modest decline in the number of quantifiable peptides over the labeling trajectory and this decline is slightly more pronounced with HW than AA labeling. Nevertheless, the results suggest that the analytical approach was able to capture isotopically labeled peptides to comparable depths in each method, regardless of the extent of labeling within the limits here. We note that HW labeling is compatible in theory with any peptide sequence, whereas AA labeling is restricted to those peptides containing that residue. When all peptides, including miscleaved peptides are admitted, regardless of the number of lysine residues in the sequence, HW labeling nonetheless quantifies approximately twice the number of peptides (**Supplemental Figure S1**).

**Figure 3.**
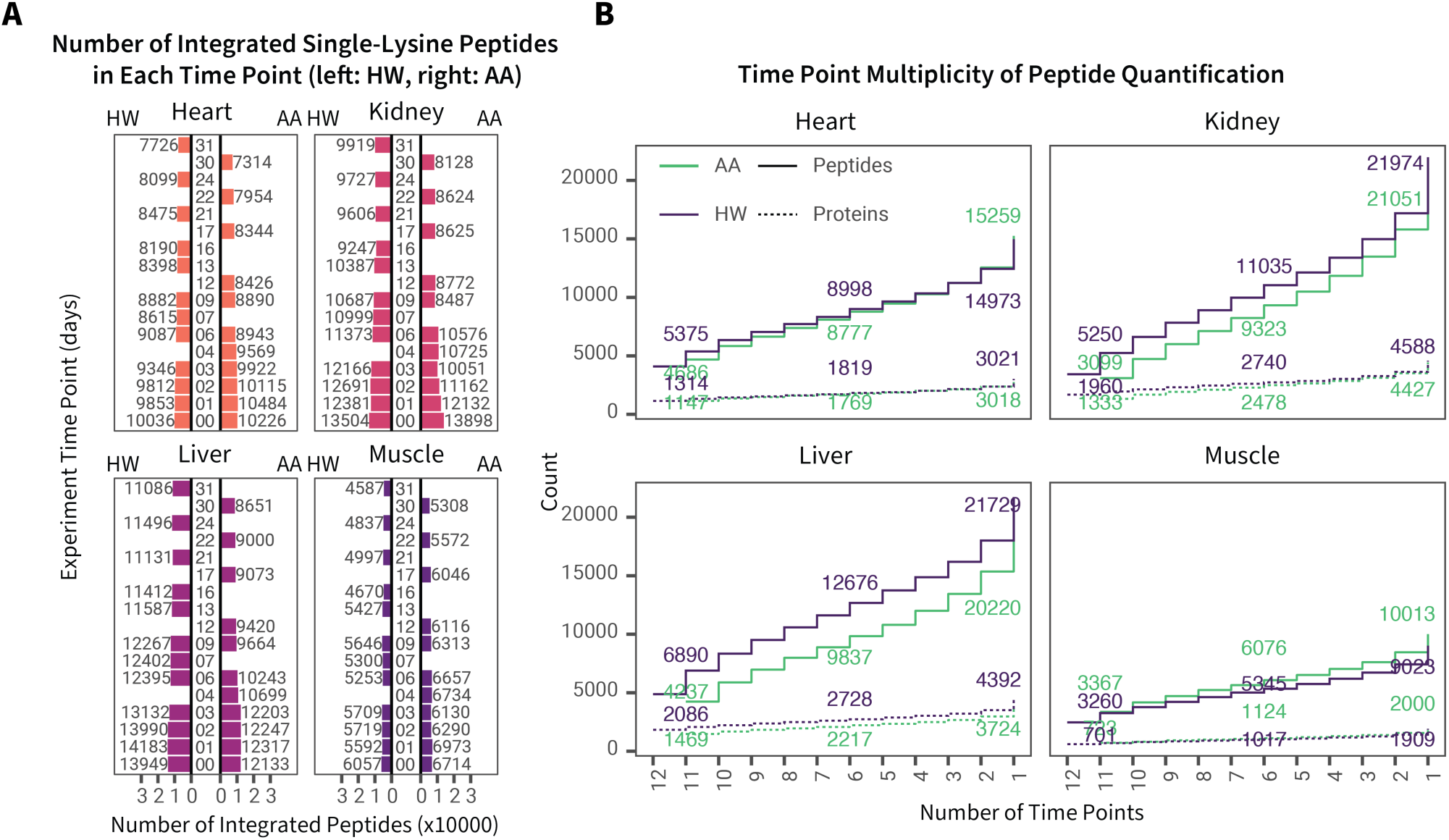
Overall depths of the comparative analysis. **A.** For each tissue, each bar defines the number of peptides integrated with Riana over each experimental time point in the labeling period. Because AA labeling requires peptides with one lysine for turnover calculation, only peptide sequences with a single lysine residue are included here for fair comparison. **B.** For each tissue, the cumulative number of peptides (solid line) and proteins (dashed lines) quantified at increasing numbers of minimal time points quantified in the heavy water (HW) labeling (green) and amino acid (AA) labeling (blue) data sets. For instance, at x=6, the y-axis numbers denote the number of peptides or proteins quantified in at least 6 time points in a labeling method and tissue.

Both labeling protocols were extended over 30 (AA) or 31 d (HW), with the first practical sampling point being at 1 d. This labeling window imposes limits on the range of degradation rate constants (*k*_deg_) that can be recovered, further confounded by the differences in rates of precursor equilibration (*k*_p_). A protein that is extensively labeled (>80%) at 1 d would have a *k*_deg_ of at least 2 d^−1^ (half life less than 8 hours). At this rate of labeling, there is no opportunity for multiple time points to define the labeling curve, and the errors in *k*_deg_ determination would be high. At the other extreme, a protein that was no more than 10% labeled at 30 d would have a *k*_deg_ of 0.003 d^−1^ or less (a half life over over two hundred days), and once again, all of the time points would have high errors, due to the low degree of incorporation. This is an inevitable consequence of stable isotope analysis by proteomic-compatible mass spectrometry and imposes analytical restrictions on the range of rate constants that can be determined.

### Analysis of raw isotopomer intensity data by non-linear curve fitting

To compare HW and AA labeling, we first used a conventional one-compartment model that is widely used in cell culture experiments *in vitro,* excluding any slow rise in labeling kinetics. The asymptotic value of the label RIA for each method can be estimated from peptides that have reached their labeling plateau, and in this study is estimated to be 0.45 for AA labels and 0.046 for HW labels. To minimize uncertainty of isotopomer quantification we used a conservative filter to admit only peptides quantified at ≥9 time points and that fitted to the one-compartment model with R^2^ of ≥ 0.9 unless specified. For the one-compartment model, the best-fit peptide *k*_deg_ values from HW and AA labeling are concordant (biweight midcorrelation ≥ ∼0.75). However, AA labeling generally reported lower peptide turnover rates compared to HW, especially apparent for peptides from proteins with relatively high turnover within a tissue (**Supplemental Figure S2**).

As stated earlier, a major difference between labeling with water and a free amino acid in the diet is the rate at which the precursor pool equilibrates. The AA data within a tissue, particularly for relatively high turnover peptides, when compared to HW labeling, suggests that AA data require a kinetic model that acknowledges this delay in equilibration in preference to a simple, one compartment model that assumes near instantaneous equilibration of label precursor pool. We therefore investigated the application of a two-compartment model to fit the HW and AA data. In the two-compartment model described by Guan et al.^23^, peptide isotope enrichment is described using two rate constants: the protein turnover rate *k*_deg_ and a composite rate constant that encompasses precursor availability kinetics; *k*_p_. This model therefore requires knowledge of the value of *k*_p_ in each tissue.

A clear indication of the behaviors of the HW and AA precursors can be gleaned from the labeling trajectories of the major urinary proteins (MUPs). MUPs are synthesized in large quantities in the liver and are immediately secreted and exported into the circulation, efficiently filtered by the glomerulus and excreted into urine, where they play multiple semiochemical roles ^44–46^. Because of the speed of this secretion and the lack of any intermediate protein pool (MUPs are very difficult to detect in plasma), the isotopomer signatures of MUPs in the liver at any time point should reflect new synthesis and rapid secretion and thus act as efficient and high speed sensors of the precursor enrichment ^47^. For HW labeled MUPs, the proteins acquire label extremely rapidly, with an average rate constant (*k*_p_) of at least 2 d^−1^ – the rapidity of labeling precludes accurate measurement of the true rate constant, but in the context of this system can be considered to be near instantaneous. By contrast, for AA labeled MUPs, the rise to plateau was notably slower, yielding a *k*_p_ of approx. 0.5 d^−1^ (half-time of about 1.4 d). If we take the AA derived value to be closest to the “ground truth” of precursor behavior, then we can compare the AA *k*_p_ rate constant with those obtained by the other approaches. Unfortunately, in the absence of true secreted proteins from other tissues that do not mix into a pre-existing pool, this insight is restricted to the liver (**Figure 4**).

**Figure 4.**
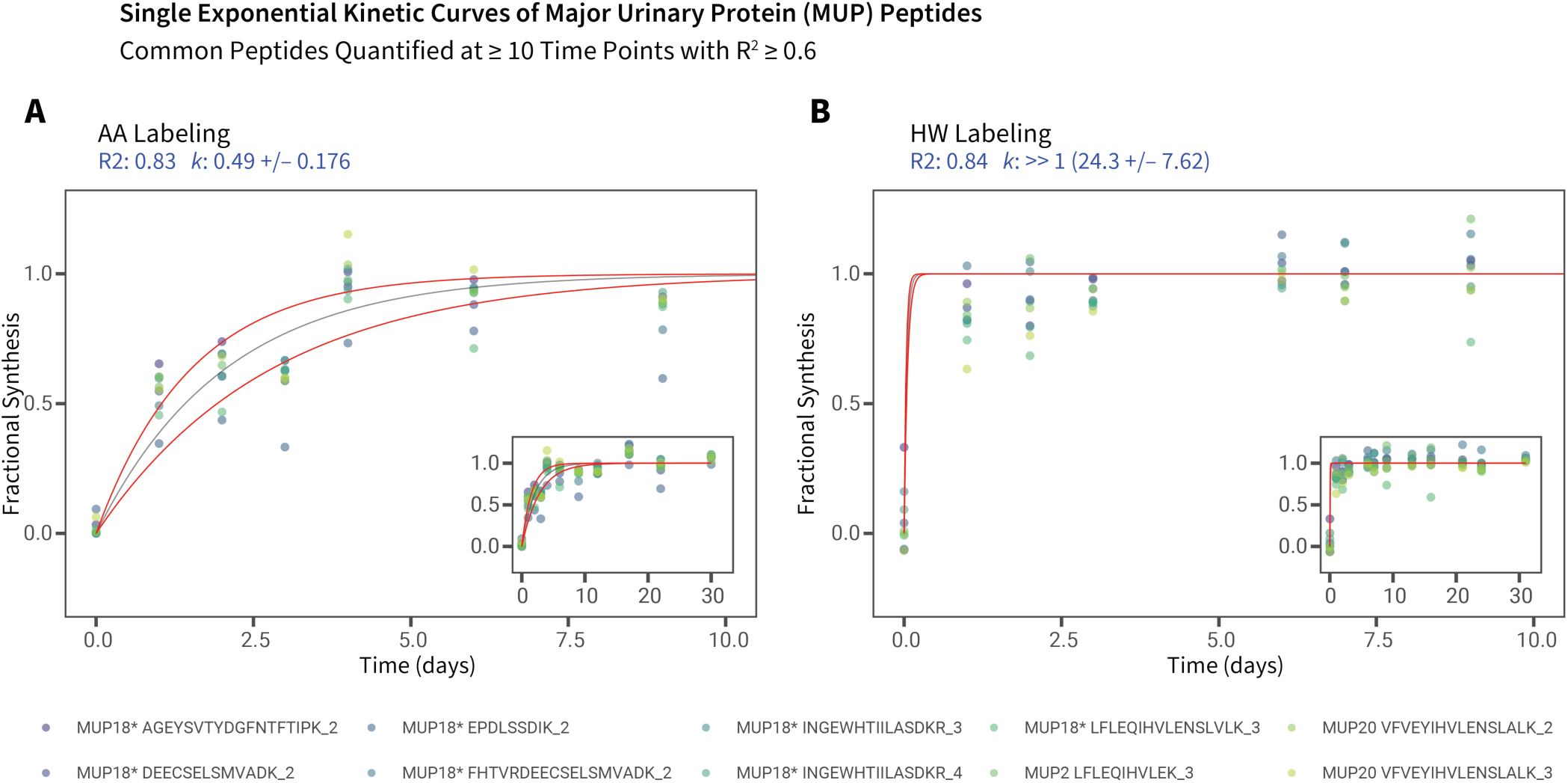
Isotopic labeling of major urinary proteins (MUPs) in the liver. Single exponential kinetic curves for major urinary protein (MUP) peptides commonly quantified at ≥ 10 time points in both the **A.** AA and **B.** HW data sets. Peptides fitted at R^2^ ≥ 0.6 at the peptide level were combined in the fractional synthesis space then fitted to a single kinetic curve to estimate the overall MUP *k*_syn_. Because MUPs are secreted from the liver as soon as they are synthesized, the quantified label trajectory is assumed to be limited only by precursor availability. As expected, in AA labeling the MUPs reflect delayed precursor kinetics with *k*_p_ of ∼0.49; whereas precursor kinetics is rapid in HW labeling with *k*_p_ >> 1. X-axis: time (days); y-axis: fractional synthesis. Main panels show expanded views from day 1 to day 10 of labeling, insets show the full data range. Asterisks after protein names denote peptide sequences present in multiple MUPs. Red lines denote models of best-fit first-order rate constant ± fitting error. The curve plotted for HW labeling cannot be construed as an accurate fit but reflects the rapidity of HW incorporation.

At the same time, because we collected both HW and AA labeling data, our experimental design allowed us to search for suitable *k*_p_ values that would bring the AA labeling data into concordance with the HW labeling data (**Figure 5A**). If we assume the HW labeling data to be more accurate for our purpose, due to the rapid precursor equilibration of water, this would suggest that a “corrected” *k*_p_ for AA labeling is 0.28–0.38 d^−1^ for heart and skeletal muscle and 0.43–0.60 d^−1^ for liver and the kidney, values substantially higher than those acquired experimentally. With these values of *k*_p_, the proteome-wide slope of log *k*_deg_ across shared peptides in HW and AA labeling approaches unity (**Figure 5B)**. Therefore, although again a two-compartment model is sufficient to describe the behavior of AA labeling data with precursor delay, it is not clear *a priori* how to produce the requisite *k*_p_ values to accurately describe tissue-specific precursor kinetics. Because the HW labeling-derived *k*_p_ values require the external reference of HW data, they are unsuitable for experiments where only AA labeling is performed. We therefore explored additional approaches that could provide a self-sufficient approach to the precursor kinetics parameters for two-compartment modeling of AA labeling. There are at least three methodological approaches by which this may be achieved: including: (1) direct empirical measurements of label in the subject; (2) a data-driven iterative approach to find the *k*_p_ that best explains the peptide data in a two-compartment model fitting; and (3) calculation of RIA using mass isotopomer distribution analysis from peptides containing two labeling sites (i.e., dilysine peptides).

**Figure 5.**
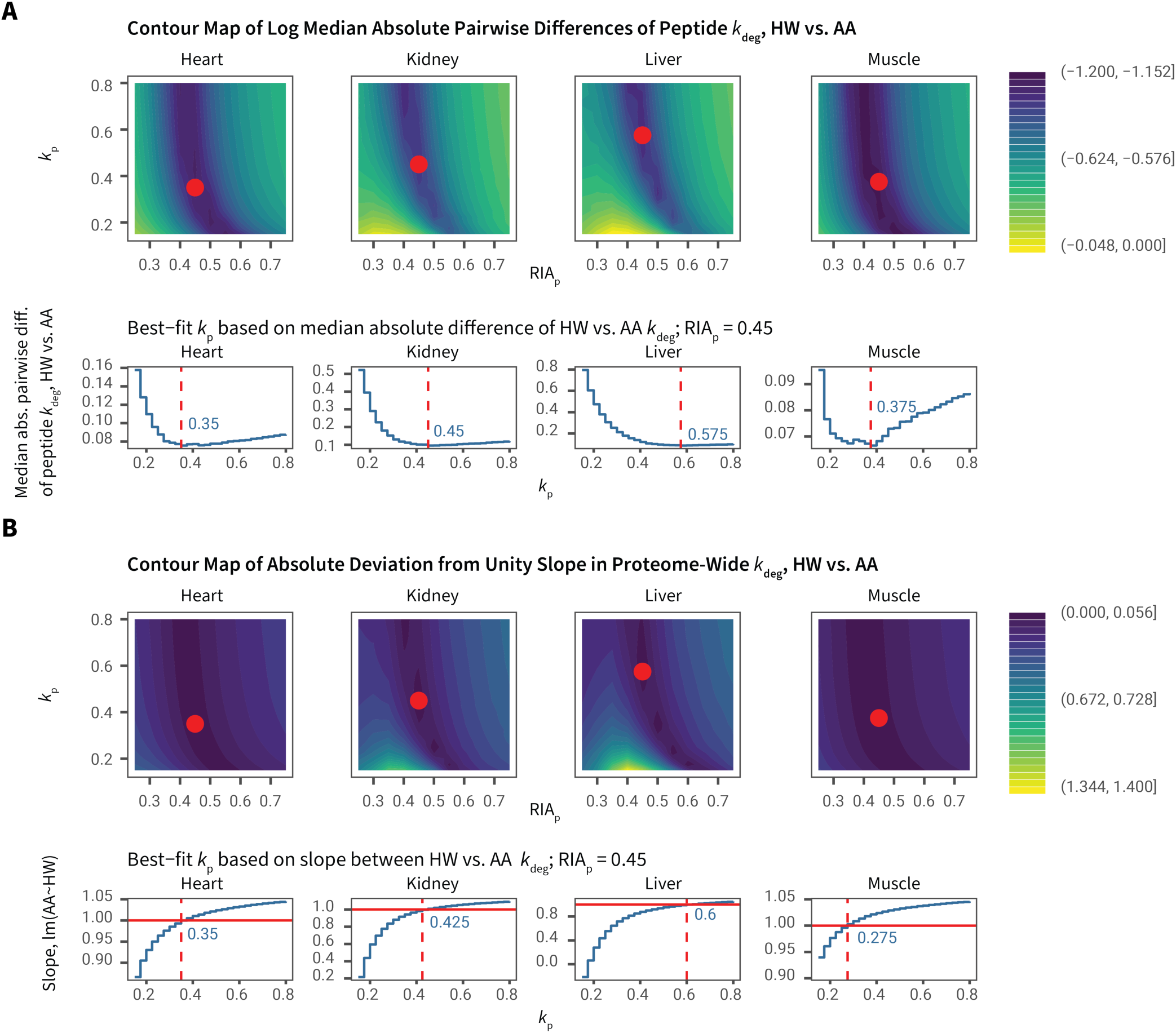
Calibrating AA labeling precursor kinetics using HW-derived rate constants. **A.** 2D density plot of the log median absolute pairwise differences in peptide *k*_deg_ from HW vs. AA labeling data, against different values of plateau RIA_p_ (x-axis) and *k*_p_ (y-axis). Red dots show the *k*_p_ values with minimal HW-AA differences at RIA_p_=0.45, also shown in the blue numbers in the line plots below. **B.** 2D density plot of the log median absolute pairwise differences in peptide-level *k*_deg_ from HW vs. AA labeling data, against different values of plateau RIA_p_ (x-axis) and *k*_p_ (y-axis). Red dots show that in the values with minimal HW-AA differences (as in panel A), the slope of proteome-wide log HW vs. AA *k*_deg_ values approach 1. Blue numbers in the line plots below show the *k*_p_ values where the slope between HW and AA *k*_deg_ are nearest to 1.

We first used LC-MS to directly measure free lysine RIA in each of the four tissues and for HW, we measured whole body water RIA enrichment by GC-MS (**Supplemental Figure S3A**). As water equilibrates rapidly across body compartments ^18^, it is assumed that the precursor kinetics for HW would be similar across tissues, but the same assumption cannot be made for AA labeling. The LC-MS determined lysine RIA enrichment curves in the four tissues vary between tissues, and did not reach a true plateau until after 30 days. This is actually inconsistent with the peptide RIA data, as the precursor enrichment cannot be slower than the peptide turnover curve. This discrepancy may be due to lysine metabolism complications or the inability to access the true protein synthesis precursor pool of lysine within the tissues (**Supplemental Figure S3B**). In fact, the LC-derived AA *k*_p_ values limit the rise of peptide RIA in the two-pool model with the consequence that the model cannot converge and does not explain the observed peptide RIA time series (**Supplemental Figure S3C**). We conclude that the LC-measured AA RIA as implemented underestimates true precursor enrichment rates and is unsuitable for explaining the peptide RIA curves and correcting for precursor delay.

We next assessed whether *k*_p_ could be gained directly from the peptide RIA data using a proteome-wide optimization approach, i.e., locate a *k*_p_ value that gives the lowest sums-of-squares (fitting error) in all peptides from kinetic curve fitting to the two-compartment model. At very high *k*_p_, the two-compartment model approaches the one-compartment model, whereas an underestimated *k*_p_ will prevent the theoretically allowable peptide (t, RIA) values from ever reaching their actual experimental measurements within the experimental time frame. Hence, the goal is to find the lowest *k*_p_ that explains the data points better in a two-compartment model compared to the one-compartment model, using the median fitting sums-of-squares of all qualifying peptides as the target. We had limited success with this nested optimization approach, and the results suggested that the strategy for finding best-fit precursor kinetics parameters will be tissue-specific. In the two low-turnover/ slow-equilibration tissues (heart and skeletal muscle), the two-compartment model outperformed the one-compartment model at a *k*_p_ of ∼0.25–0.35 d^−1^. The two-compartment model never fitted the data better than the one-compartment model in the liver nor the kidney (**Supplemental Figure S4A–B**). On the other hand, proteome-wide optimization over the fast-equilibration tissues allowed the effective plateau precursor RIA (RIA_p_) in these tissues to be estimated directly from the data, which was not possible in the slow-turnover tissues. We interpret the results to suggest that although a two-compartment model is necessary to correct for labeling delay, the precise values of *k*_p_ cannot be easily found in fast equilibration tissues as different combinations of *k*_p_ and *k*_deg_ yield identical kinetic curves. Conversely, in some tissues, the best-fit *k*_p_ learned from the data will not necessarily give accurate absolute values of peptide *k*_deg_.

Finally, we estimated precursor RIA over time using mass isotopomer analysis with dilysine peptides. This is a commonly used method in dynamic SILAC studies in animals, where the heavy-heavy and heavy-light peaks of a peptide containing two labeled amino acids is used to reveal the true precursor RIA during the time when the peptides were made ^5^. However, there is no commonly accepted standard for selection of the dilysine peptide(s) for this calculation. We therefore calculated the precursor RIA, restricted to all dilysine peptides quantified at a minimum of 9 time points in each tissue. Whilst there is a noticeable increase in estimated precursor RIA as labeling proceeds (**Supplemental Figure S5A**), the calculated precursor RIA values from each peptide has high variance, especially at earlier time points. Using a Gaussian kernel density estimate, we estimated the mode RIA at each time point as the representative tissue precursor RIA (**Supplemental Figure S5A**). The resulting tissue RIA estimates fitted well to single exponential curves. However, the derived *k*_p_ values remain lower than required to explain the peptide curves in AA labeling or the MUP-derived prediction, and the curve fitting is unable to converge to a satisfactory solution for a number of peptides.

As an alternative to kernel density estimates, we used all qualifying tissue-wide RIA_p_ values (dilysine peptides identified at ≥9 time points, 0 ≤ RIA ≤ 0.6) to define a single exponential kinetics model using weighted nonlinear least square fitting across each tissue to estimate the precursor rate constant and plateau (**Supplemental Figure S6**). The contribution of each RIA data point to the fitted curve is weighted by the square of the peptide isotopomer normalized intensity. This approach produces *k*_p_ values that align well with those derived from HW adjustment. For the liver, the MUP-derived *k*_p_ was essentially the same as the dilysine recovered parameter (0.49, *cf.* 0.52 d^−1^)

Because the RIA curve in the weighted fitting exhibited biphasic behavior, we also fitted the tissue RIA curves to the two-exponent kinetic model described by Fornasiero *et al.*^11^, which accounts for label dilution from global protein degradation, using three parameters, the soluble precursor enrichment/breakdown rate constant, *b*, the global protein degradation rate constant, *a*, and the ratio of lysine pool in soluble vs. protein-bound pools in a tissue, *r* (**Supplemental Figure S6**). Reutilization, represented by *a,* contributes to the slow phase precursor rise following the initial plateau as the reutilized lysine residues originating from protein degradation products slowly become labeled. The two-exponent kinetic model fitted to dilysine peptides derived RIA values significantly better than the single exponent model, accounting for the extra number of parameters (Akaike weights 1 to 9e–44 in the heart; 1 to 1e–27 in the kidney; 1 to 1e–27 in the liver; 1 to 9e–18 in the muscle). When incorporated into a two-compartment, three-exponent model (*a*, *b*, *k*_deg_) for individual protein *k*_deg_ values, the Fornasiero *et al*.^11^ method yielded results comparable to the two-compartment, two-exponent model in Guan et al.^23^ for slow-turnover peptides within a tissue, but was able to correct for fast turnover proteins within a tissue in the AA labeling experiments (**Supplemental Figure 7A–B**). Both two-compartment models led to higher intra-protein variance (**Supplemental Figure 7C**) than the single exponential model (see paragraphs below) across various R^2^ cutoffs. The different methodologies and resultant values for various precursor kinetics parameters are summarized in **Table 1**.

**Table 1.**
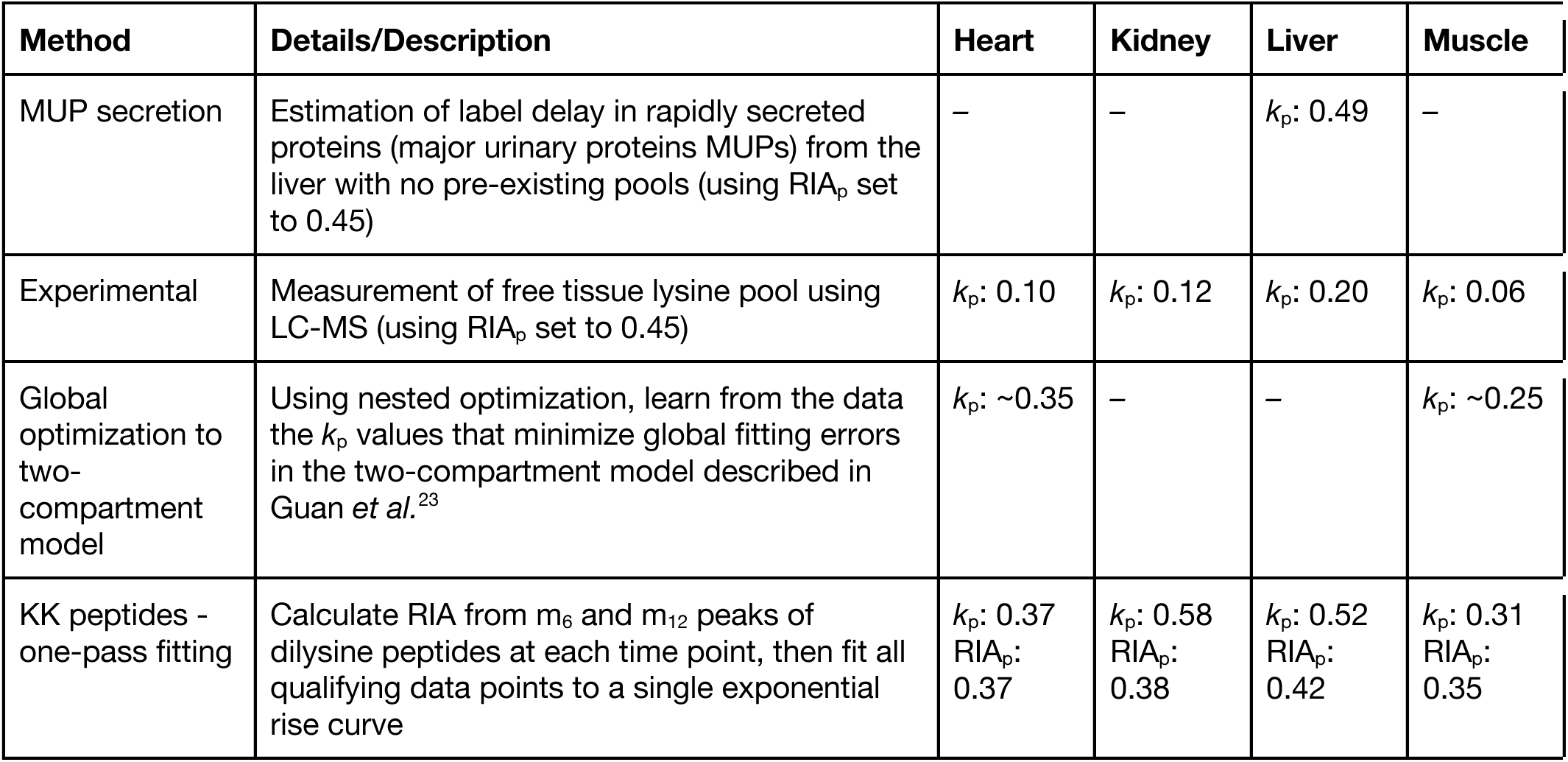
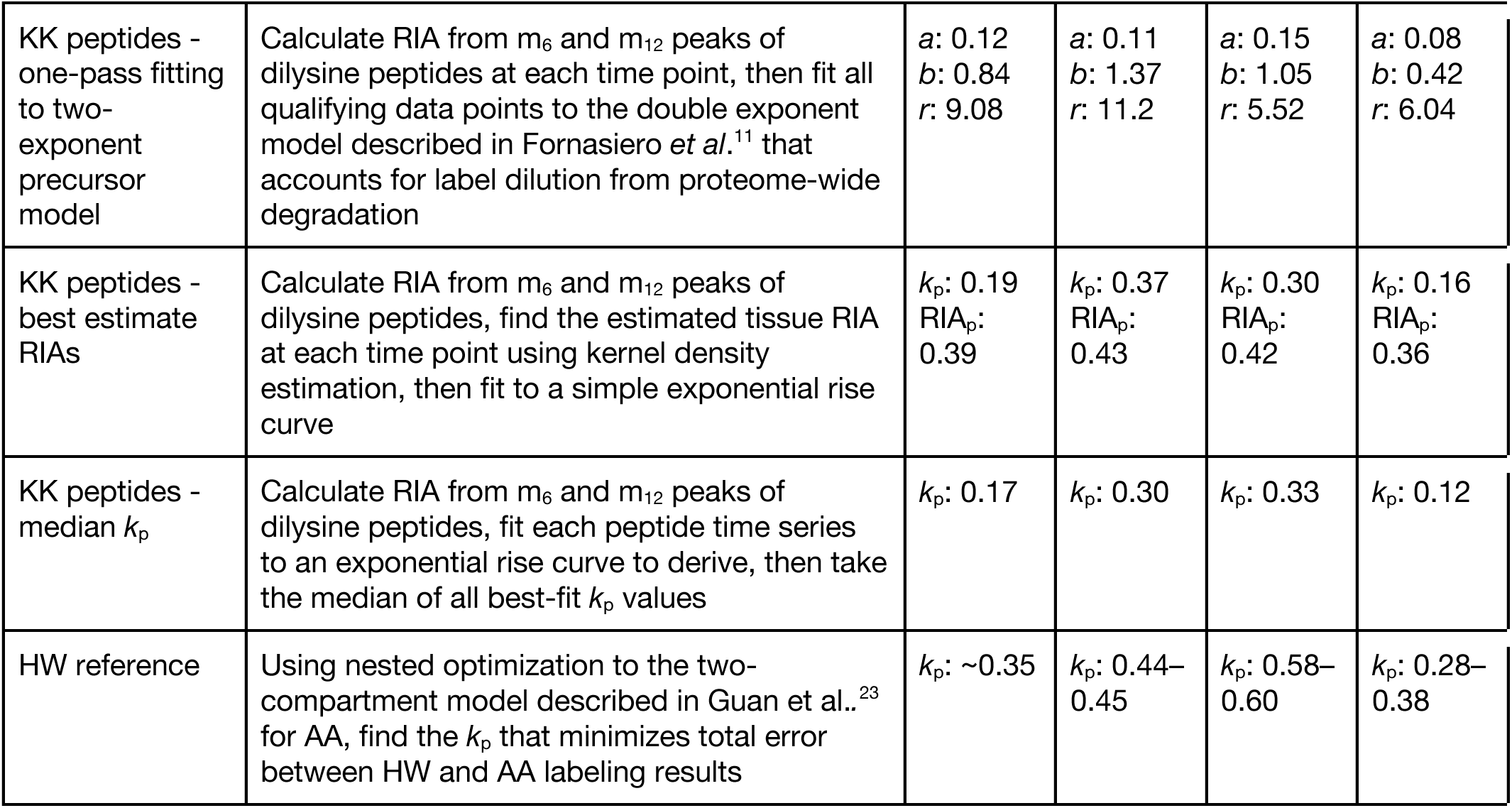
Methods for determination of AA labeling precursor kinetics. *k*_p_: precursor relative isotope abundance kinetics rate constant; RIA_p_: asymptotic precursor relative isotope abundance.

From these analyses, and using the MUP-derived parameters as ground truth, we conclude that correction for slow equilibration (whether caused by slow uptake or reutilisation) is feasible, and that analysis of dilysine peptides, two-pool modelling or reutilisation correction give the best estimates.

### Comparison of peptide turnover rates between HW and AA labeling

We next used the weighted dilysine peptide *k*_p_ and asymptotic RIA_p_ values in a two-compartment, two-exponent model to derive peptide *k*_deg_ for all qualifying single lysine peptides in each tissue. We were thus able to fit the AA (slow equilibration) and HW data (rapid equilibration) using either model for comparison. We compared the best-fit peptide *k*_deg_ from the one-compartment model to a two-compartment model that accounts for the delay in precursor equilibration. As anticipated, the two-compartment model influenced the kinetics of the AA labeling experiment more than HW labeling, particularly for higher turnover proteins within a tissue. This is evident from the off-diagonal distribution between one-compartment and two-compartment models in AA labeling as well as the absolute differences in high-turnover peptides (**Figure 6A**).

**Figure 6.**
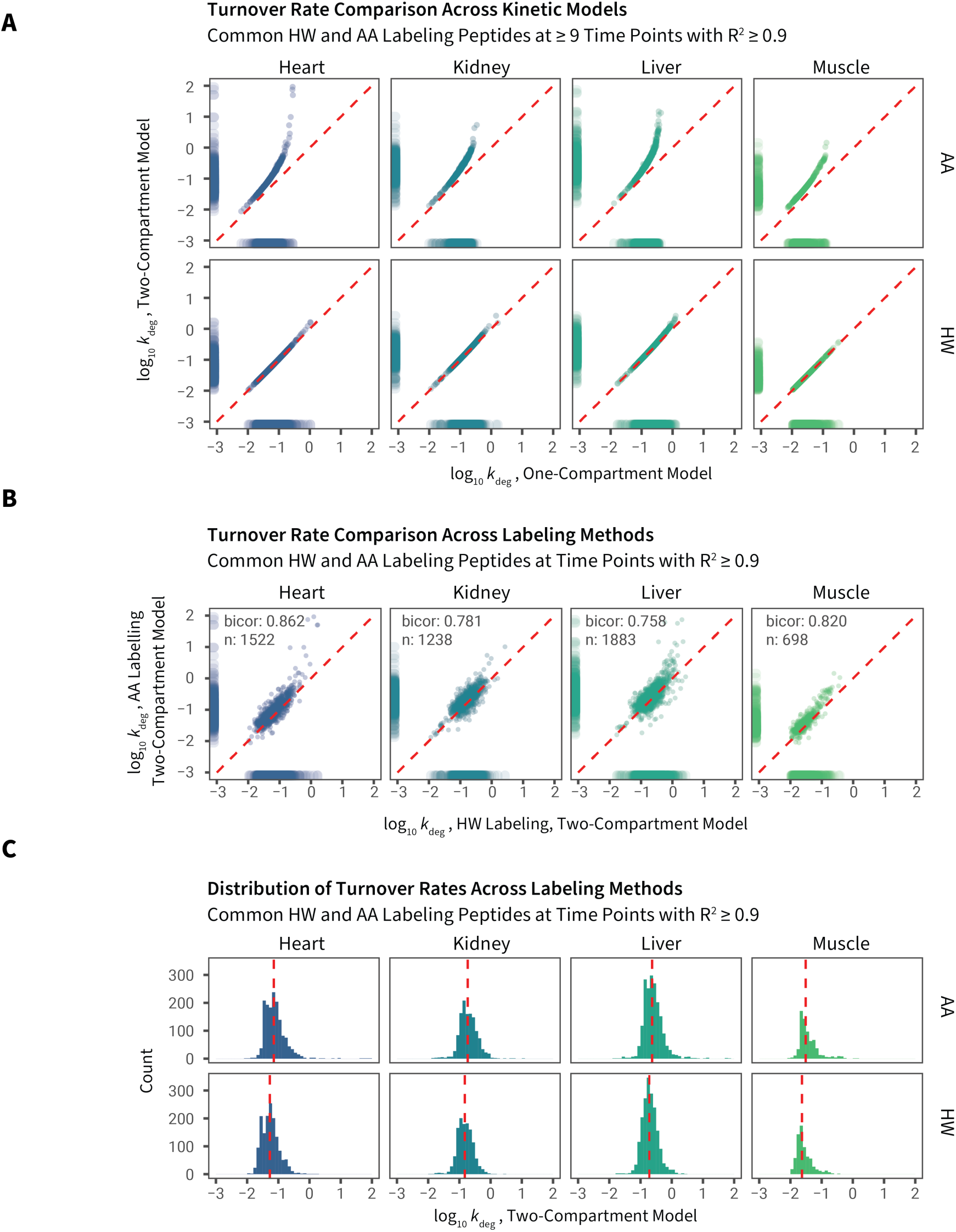
Comparisons of turnover rate constants across labels and kinetic models. **A.** Degradation rate constants (*k*_deg_) were obtained for proteins from four tissues using amino acid (AA) or heavy water (HW) labeling, and were fitted using a one-compartment (x-axis) or two-compartment (y-axis) kinetic model to derive the first order *k*_deg_ (plotted on a log base 10 scale). Data points are peptide time series with ≥ 9 time points and fitted with R^2^ ≥ 0.9. Red dashed line: unity. Marginal distribution shows data density. **B.** Scatterplots of shared proteins quantified by HW and AA in each tissue using the two-compartment model (quantified time points ≥ 9, R^2^ ≥ 0.9). Numbers denote robust correlation (biweight midcorrelation; bicor) coefficients. Each data point is one peptide-charge time series. **C.** Histogram showing distribution of *k*_deg_ across tissues and labels. Red dashed lines denote medians.

Fitting to the two-compartment model brought the median turnover rates of peptides quantified from the two methods into closer correspondence (**Figure 6B**), and corrected the discrepancy between HW and AA labeling in relatively high turnover proteins within a tissue (**Figure 6C**). For peptides integrated over at least 9 time points and fitted with an R^2^ of ≥ 0.9, the agreement between the methods is good (robust correlation bicor: 0.758–0.862 in four tissues) when the two-compartment model is used to fit the (peptide RIA,t) data. Both AA and HW labeling data showed excellent agreement with the turnover rates derived in a previous study of HW labeling of heart proteins an independent cohort of C57BL/6J mice ^15^ (biweight midcorrelation: 0.85–0.93; **Supplemental Figure S8A**) and good agreement with a more recent study of AA labeling of liver and skeletal muscle proteins in NSBGW mice ^24^ (biweight midcorrelation: 0.73–0.83; **Supplemental Figure S8B–C**). The *k*_deg_ values of peptides in HW and AA labeling are tabulated in **Supplemental Data S1**. All fitted peptide curves using the two-compartment model above in AA and HW labeling are in **Supplemental Data S2–S5**.

A few peptides showed unexpectedly high turnover rates in AA labeling, possibly because the precursor enrichment rate is underestimated, leading to overcorrection. For these peptides it is likely that *k*_deg_ could not be determined with high accuracy as the non-linear model would fail to converge when it is limited by *k*_p_, and these peptides may be more accurately categorized simply as having high turnover. Nevertheless, we conclude that the two-compartment model performed well in correcting the underestimation of *k*_deg_ for relatively high-turnover proteins (*k*_deg_ ≥ 1 d^-1^); proteins for which label enrichment is retarded measurably by the lag in precursor enrichment (∼1 d^-1^). This is more pronounced in tissues where the overall protein turnover rates are high, such as the kidney and the liver.

### Evaluation of data quality filters

Using the one and two-compartment model results, we examined the effect of various data quality filters on the depth and reliability of protein turnover measurements. We varied the filtering criteria by two metrics: first, the number of time points in which the degree of label incorporation into protein was quantifiable; and secondly, the coefficient-of-determination (R^2^) of the one-compartment and two-compartment kinetic models. We then considered their effects on interaction with the number of quantifiable peptides, and the precision of turnover rates. To estimate precision, we considered the median values of the geometric coefficient of variation (CV) in turnover rates among peptides uniquely mapping to the cognate protein, because peptides derived from the same protein should be synthesized and degraded together *in vivo.* Barring undocumented proteoforms, peptides from the same protein should yield identical turnover rates, if the measurement is precise.

First, we observed an expected decrease in the number of available peptides as the thresholds for required time points and R^2^ were raised. There was a sharp decrease in quantifiable peptides at all time point cutoffs when the R^2^ threshold increased beyond ∼0.8, suggesting a drastic decrease in profiling depth if too stringent a threshold is used (**Figure 7A**). Although the number of quantifiable peptides between the one-compartment and two-compartment models at each R^2^ threshold are similar, there is a noticeable difference in intra-protein geometric CV. We found that the two-compartment model led to increased variance at all R^2^ cutoffs. This increase was especially noticeable in fast equilibration tissues (liver and kidney), and is especially severe in AA labeling (**Figure 7B**). This is probably attributable to the two-compartment model having more parameters that can vary, e.g., true *k*_p_ may differ across cell types within a tissue, demanding more stringent time-point and R^2^ thresholds. In our experience, data quality and the ability to make inference about changes in turnover declined when the median geometric CV increased beyond ∼0.33.

**Figure 7.**
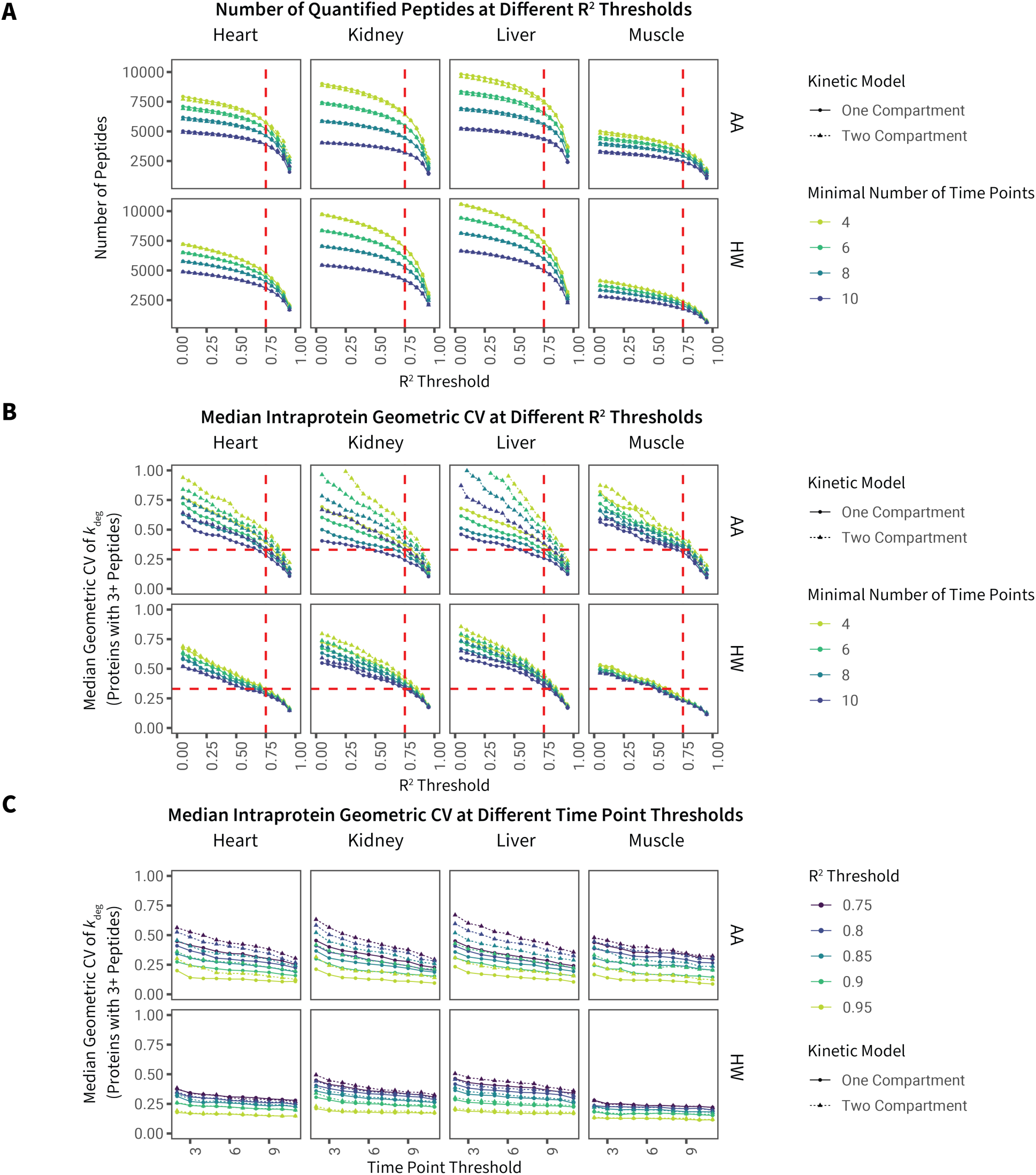
Relationship between R^2^ and time point filters on peptide count and variance. **A.** The number of quantified peptides (y-axis) vs. R^2^ coefficient-of-determination thresholds in kinetic curve-fitting (x-axis) with various time point filters (colors). Both R^2^ and minimal time points have a large effect on the total number of fitted peptides. Red dash lines: R^2^ 0.75. **B.** Intra-protein variance, measured as the geometric coefficient of variation (CV) of best-fit *k*_deg_ among peptides uniquely mapped to the same proteins (y-axis), vs. R^2^ thresholds (x-axis) and time point thresholds (color). Only peptides belonging to proteins with 3 or more quantified peptides were used for the analysis. Two-compartment models led to higher intra-protein variance. Horizontal red dash lines: geometric CV = 0.33; vertical red dash lines: R^2^ 0.75. **C.** Intra-protein variance (y-axis) as in panel B, against the minimal number of required time points, at different R^2^ thresholds. It can be seen that R^2^ has a more pronounced effect on *k*_deg_ precision than minimal time point thresholds.

When an R^2^ threshold of 0.8–0.95 is imposed, the inclusion of peptides quantified at fewer time points had a modest impact on intra-protein *k*_deg_ variance (**Figure 7C**) while allowing more quantifiable peptides (**Figure 7A**). Overall, we surmise that with a two-compartment model, a conservative R^2^ threshold is needed to minimize intra-protein variance whereas the number of quantifiable data points has a lesser impact. Hence, we selected an R^2^ threshold of 0.9 and a time point threshold of 6 to investigate proteome level features of protein turnover in the four tissues.

### Protein-level data summary

Finally, we aggregated peptide-level turnover rates to the protein level, by selecting well-fitted peptides (R^2^ ≥ 0.9, 6 time points) and refitting all fractional synthesis data points to a single curve. We found little difference between an unweighted fitting and fitting weighted by normalized log peptide intensities. The *k*_deg_ values of proteins are tabulated in **Supplemental Data S6**. All individual protein-level kinetics curves using the two-compartment model above in AA and HW labeling are at **Supplemental Data S7–S10.**

## Discussion

There is a need for rigorous assessment of the strategies used to measure protein turnover rates in intact animals. Irrespective of the labeling precursor, there is potential for a delay in the equilibration of the precursor pool. Although this can be ameliorated by the use of HW as a precursor label, knowing the precursor enrichment kinetic rate constant (*k*_p_) in animal studies is recognized as a challenge. The primary focus of our study is the comparison of two labeling methods (HW vs. AA) with different precursor equilibration rates across high- and low-turnover tissues. We also explore different approaches to derive the precursor RIA kinetics parameters in AA labeling, but none of the three approaches taken is entirely satisfactory for all tissues. If we assume the HW fitted data have higher reliability due to the minor impact of complications from precursor delay, then tissue-specific values of the precursor enrichment rate constants allow the AA data to align closely with the HW data. These values were ∼0.28–0.35 d^−1^ for the adult mouse heart and muscle and ∼0.44–0.6 d^−1^ for the liver and kidney, but it was not entirely clear how they might be reliably derived without the external reference of HW data.

Surprisingly, direct LC-MS measurement of intracellular lysine pools gave *k*_p_ values that were considerably lower than required to be compatible with label enrichment of peptides. There may be preferential label reutilisation of amino acids released by degradation for synthesis *de novo*, such that the measured free lysine is decoupled from the true precursor pool. Other complications may also cause the precursor enrichment rate to deviate from exponential rise kinetics, for instance, unlabeled lysine is dietary and delayed by digestion prior to transport, whereas labeled lysine is a free amino acid that is available to membrane transporters immediately. Analytical approaches might require complex measurements of precursor RIA in multiple “compartments”: plasma, interstitial fluid, intracellular fluid as well as in the aminoacyl tRNA pool.

We therefore explored two alternatives to estimate *k*_p_, but neither approach was entirely satisfactory. We examined a two-compartment nested optimization method to gain the best fit *k*_p_ that explains the labeling curves directly from the peptide data. This method may compensate for the time integral in the dilysine peptide analysis, and indeed appears to perform well for slow-equilibration tissues such as the heart and the skeletal muscle. However, when *k*_deg_ << *k*_p_ for most peptides in a fast-equilibration tissue such as the liver, the two-compartment model never outperformed the one-compartment model in minimising fitting errors, likely because the initial sigmoidal “bend” in the kinetic curve of the two-compartment model is not apparent and thus different combinatorial values of *k*_p_ and *k*_deg_ can explain the peptide RIA data equally well.

We have previously advocated an approach based on mass isotopomer analysis of peptides containing two instances of the amino acid ^5^. This approach has the potential to reveal the true precursor isotopic enrichment based on the immediate protein precursor pool (e.g., labeled aminoacyl tRNA), but there is a technical challenge associated with determination of precursor pool behavior. Because all subsequent protein turnover measurements use these kinetics that define precursor behavior, it is important to analyse doubly-labeled peptides carefully, either by manually extracting clean and readily isolatable extracted ion MS1 chromatograms, or by automating analysis of all peptides and modeling the best estimates for each time point. Unexpectedly, there is a wide distribution of calculated RIA values from HH and HL peaks. A best-fit exponential model of RIA over time has large residuals but yields values of *k*_p_ consistent with expectation. The RIA fitting also shows distinct best-fit RIA_p_ plateaus in each tissue, which is counterintuitive - if the subject had been exposed to the labeled AA during uterine development and at all times subsequently, the RIA_p_ for each tissue should be identical. Over the 30 d labeling window used here, the precursor pool would be diluted by unlabeled lysine derived from the pre-existing protein pool (the subjects are adult, non-growing). Further, this approach will suffer from coupling to the rate of turnover of the protein when *k*_p_ is not constant or the protein has not completely turned over. A very high turnover protein (one that might not be measurable with any accuracy, as tissues cannot be sampled rapidly enough) would be replaced quickly, and thus become a high frequency sensor of the precursor pool. By contrast, a protein with a lower rate of replacement would retain a significant proportion of the protein pool that was synthesized in the early stages of precursor pool equilibration, thus giving a dampened measure of the rate of precursor rise to plateau, although the plateau value for RIA should be the same in either instance. However, empirically the *k*_p_ values calculated from dilysine peptides are not correlated with the *k*_deg_ of the peptides.

These complications serve to highlight the difficulties of using labeled amino acids in intact animal systems. All of the solutions that have been explored here are complicated and require additional analyses or complex modeling. Even direct measurement of the tissue free amino acid pool might not be suitable or representative of the true precursor of protein synthesis; the aminoacyl tRNA pool. Optimisation of complete data sets to derive precursor behavior may introduce uncertainty, and the error gradients are shallow for many combinations of *k*_p_ and *k*_deg_.

Taken together, we proceeded with the *k*_p_ values from automated dilysine analysis as they led to *k*_deg_ that agreed with HW data. A more complex two-compartment model ^11^ that accounts for label dilution from global protein degradation further improved data fit and concordance with HW results incrementally. However, one is left with the same problem of having to learn the model parameters from the data without guarantee that such values can be found. Moreover, even if a complex model fits the data more closely, it cannot be guaranteed that the resulting optimized *k*_deg_ are in fact accurate values as they are inside the cell. Prior studies that compared fitting models largely used only two criteria to compare different models - (i) measures of model fit such as residual sums of square, coefficients of determination, or information criteria; (ii) the ability of the model to avoid unreasonable kinetic rate constants in the fitted results, e.g., negative values, or values that are out of the allowable numerical ranges given the sampling time points. Generally speaking, however, the introduction of additional parameters in a complex model will allow the kinetics function to move more freely to the data points, which can lead to trade-offs between variance and bias and the possibility of overfitting.

At present, we are unaware of an accepted gold standard of turnover rates of proteins *in vivo*. Although premixed synthetic isotope analogs with known isotope ratios may be used as a standard for the accuracy of mass spectrometry quantification of isotopomer intensities, they cannot serve as a calibration target of *in vivo* turnover rates or precursor enrichment. Literature values of turnover rates *in vivo* remain few, and turnover rates vary greatly by species, tissues, age, and physiological states, making direct comparisons difficult. In the absence of *k*_deg_ standards, we propose that intra-protein variance should be examined as a criterion by which analytical methods to derive turnover rates are evaluated. This is based on the simple assumption that a protein is created and destroyed in its entirety (i.e., a protein is never partially excised from one terminus then repaired with replacement amino acids). Hence different peptides from the same protein should share similar turnover rates, if the modeling results are reliable, and by extension, this gives an estimate of whether the model will return identical *k*_deg_ for two polypeptide chains with equal true turnover rates. We found that the two-compartment model generally increases intra-protein variance at equal R^2^ cutoffs, hence stringent data filtering strategies are needed for the two compartment model fitting in order to maximize the number of quantified peptides while minimizing intra-protein variance.

For robust measurement of turnover rates in intact animals, we would recommend: (a) labeling with [^2^H_2_]O, (b) determination of labeling profiles in a peptide-specific analytical workflow to compensate for the specificity of HW labeling, (c) distribution of replicates in the time domain, to define as broad a range of turnover rates as possible, (d) aggregation of data from multiple peptides to increase confidence in the extraction of the MS1 isotopomer profiles, (e) stringent quality filters at the peptide level in the analysis methods to minimize intra-protein variance. In reporting turnover rates, care should be given to the limitations imposed by the sampling time points - these intervals set limits on the range of turnover rates that are accessible. This is eased by a temporally expanding sampling window. For example, sampling of 1, 2, 3, 4, 8, 10, 13, 16, 20, 32 d, and working on the assumption of a measurable abundance for labeled or unlabeled peptide of 5% of the total, over three data points, should yield measurable variation in the RIA for proteins with k_deg_ values between 0.002 d^-1^ and 1 d^-1^. Beyond these limits is it only possible to say ‘less than 0.002 d^-1^’ and ‘greater than 1 d^-1^’. Ideally, sampling intervals less than 1 day would allow access to higher turnover proteins, arguably only feasible with HW strategies. At the other extreme, very low turnover proteins could require labeling periods of many months, with less frequent sampling.

This study is a highly controlled comparison of HW and AA labeling strategies for measurement of peptide turnover rates in four tissues in intact animals. In this paper we have evaluated strategies for high quality measurements of turnover rates - discussion of the biological significance of these results will be addressed in a separate publication. It is possible to bring the two data sets into close agreement when the AA precursor behavior is addressed to provide a suitable *k*_p_ correction. AA labeling is analytically simple but metabolically complex, with tissue-dependent variation in behavior. Given the challenge of determination of *k*_p_ values for a two-compartment model in AA labeling, dilysine peptides and iterative two-compartment model fitting performed better than direct measurement of free lysine in tissues. These approaches provided *k*_p_ values that are consistent with the observed peptide data. Improved methods to model the RIA values of double-labeled peptides, such as a time integral of protein pool replacement, could improve measurements using AA labeling approaches. By contrast, HW equilibrates quickly and non-enzymatically and hence *k*_p_ is systems-wide across tissues, and requires minimal correction in precursor pool equilibration. HW labeling is also cheaper and enables the quantification of a substantially greater number of peptides. However, isolation and analysis of the subtly shifting isotopomer profile is more challenging.

Future studies of whole animal protein turnover will require even greater attention to a range of prerequisites, including choice of label - at present, the strongest case is for the use of heavy water. Further, the duration of the labeling experiment (to cover the broadest range of turnover rates that are required) is critical, although it will always be challenging to recover accurate rate constants for particularly high and low turnover proteins. There is also a case for an increased number of time points to scribe the labeling curve - we would argue that points distributed across the labeling curve are preferable to replicate samples with fewer time points. Lastly, there is a case for a well conducted labeling study to be analysed by many of the analytical packages for turnover determination - a detailed comparison of the resultant output would be especially informative. This would lead us to a more open democratisation of the analytical workflow, and a clearer understanding of where there are variances in the final outputs. To this end, all raw data from this study are available for immediate download. Lastly, an animal that has been labeled in such studies is often the source of just one or a few tissues. The ‘3R’ principles underpinning animal research (Replacement, Reduction, Refinement), specifically, reduction, would be well served if, in future, labeling and turnover studies were enhanced by a willingness to share unused tissues, to allow others to replicate studies, improve data analysis and extend understanding. Indeed, in a recent review a future imperative was exactly this; “Data and tissue-sharing offer opportunities for more efficient use of information collected from animals and may avoid unnecessary repetition” ^48^.

## Acknowledgments

This work was supported in part by grants from the BBSRC (BB/J002631/1) and The Leverhulme Trust (RPG-2019-192) to JLH and RJB; and US NIH/NHLBI Funding R00-HL144829 to EL.

## Supplemental Figures

**Supp. Figure S1.**
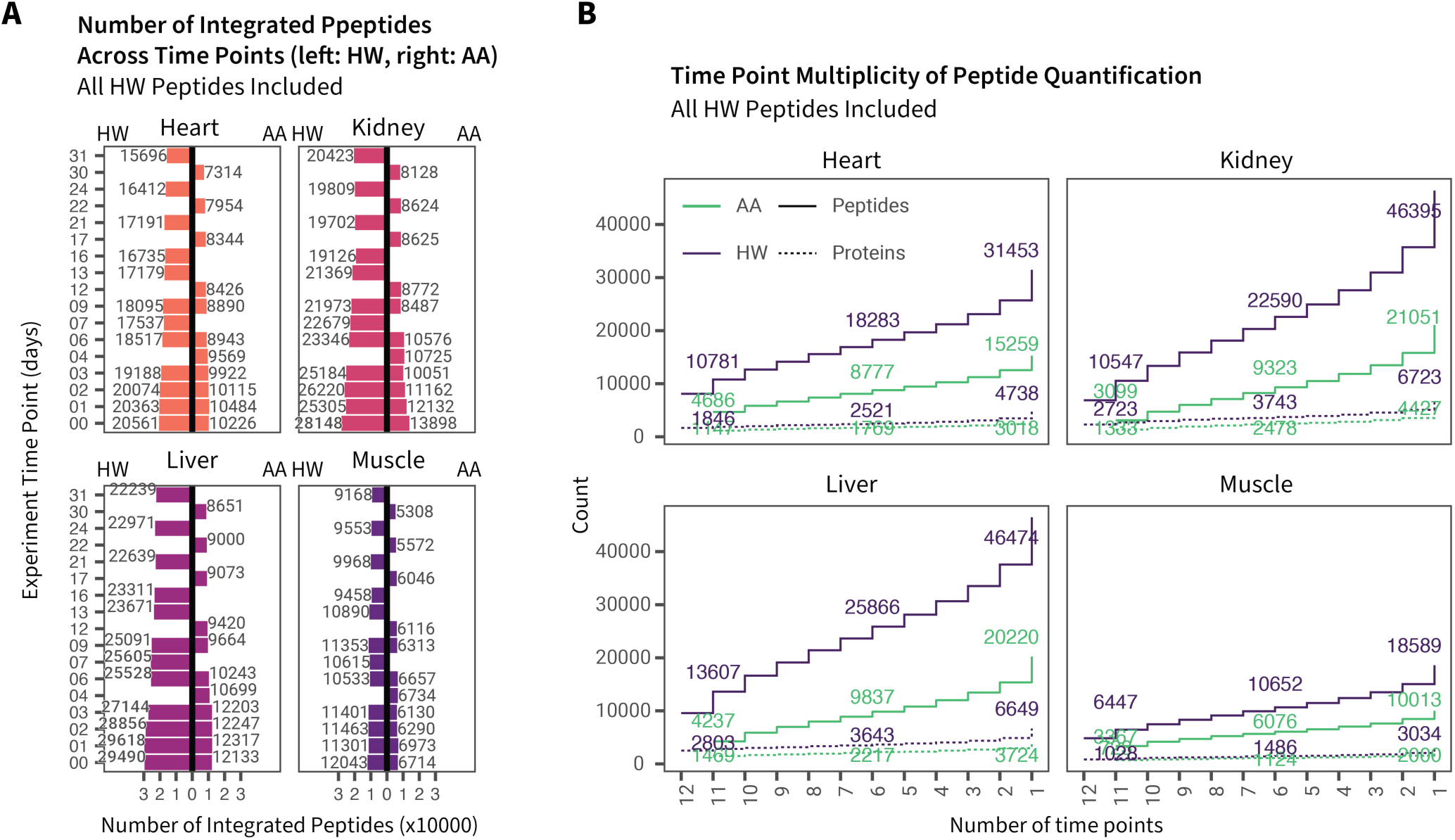
Peptide quantification performance inclusive of all HW labeling peptides. **A.** As in Figure 3, for each tissue, each bar defines the number of peptides integrated over each experimental time point in the labeling period. Here only peptides containing one lysine are included in the AA labeling experiment, whereas all quantified peptides are included in the HW labeling experiment. **B.** For each tissue, the cumulative number of peptides (solid line) and unique proteins (dashed lines) quantified at increasing number of minimal time points in the heavy water (HW) labeling (green) and amino acid (AA) labeling (blue) data sets.

**Supp. Figure S2.**
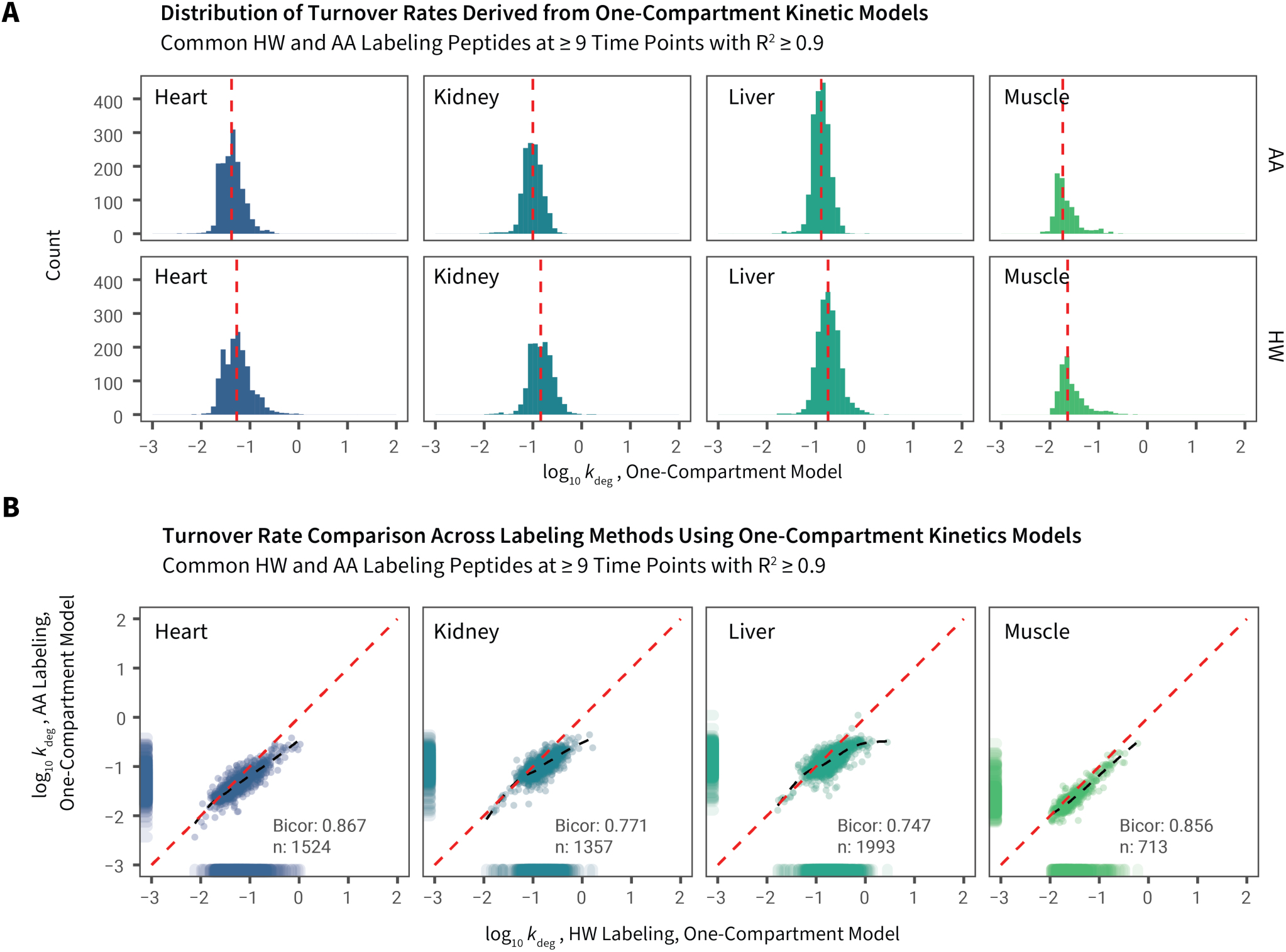
Comparison of HW and AA labeling data in one-compartment fitting. **A.** Histogram showing distribution of *k*_deg_ across tissues and between HW and AA labels using a simple exponential model (quantified time points ≥ 9, R^2^ ≥ 0.9) where peptide isotope enrichment is described by a single rate constant (*k*_deg_). Red dashed lines denote medians. **B.** Scatterplots of shared proteins quantified by HW and AA in each tissue using the one-compartment model (quantified time points ≥ 9, R^2^ ≥ 0.9). Numbers denote robust correlation (biweight midcorrelation; bicor) coefficients and numbers of compared peptides (n).

**Supp. Figure S3.**
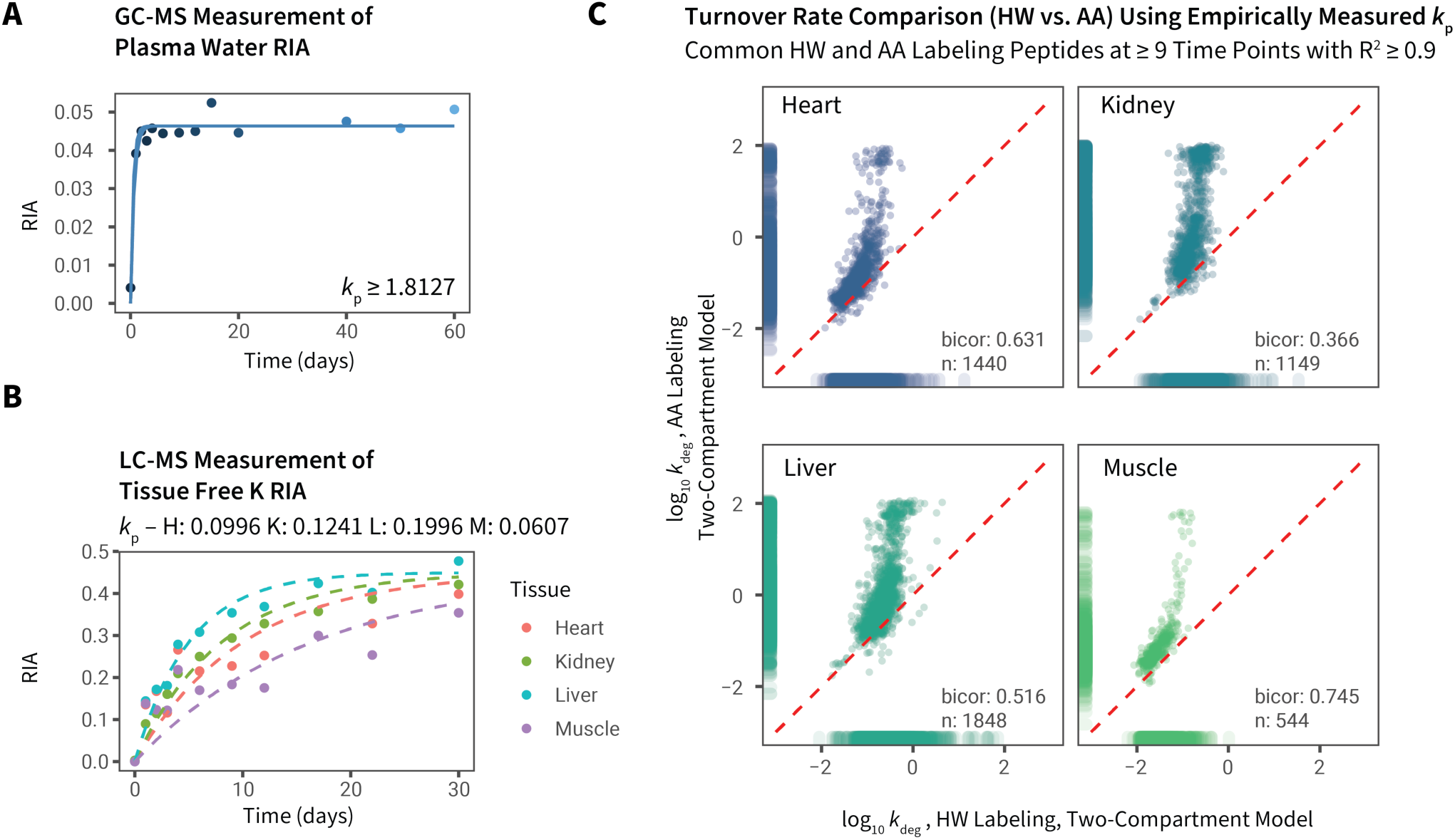
Empirical measures of tissue precursor RIA values at each time point. Precursor relative isotope abundance (RIA_p_) was measured using **A.** GC-MS of plasma samples in HW labeling and **B.** LC-MS of tissue free lysine in AA labeling. The precursor RIA over time data were fitted to a simple exponential model to find the best fit *k*_p_ using nonlinear least squares. **C.** Using the empirically-derived *k*_p_ values in a two-compartment model led to apparent high *k*_deg_ peptides in AA labeling, since the peptide RIA rise curves for fast-turnover peptides do not converge to the model when it is constrained by an underestimated *k*_p_. X-axis: log_10_ turnover rate constants (*k*_deg_) of HW labeling; y-axis: log_10_ turnover rate constants of AA labeling. Each data point represents one peptide. Peptides integrated at ≥ 9 time points and fitted to a two-compartment model at R^2^ ≥ 0.9 are included. Red dash line: unity. Number: biweight midcorrelation (bicor) and number of individual peptides compared in HW vs. AA. In this panel, the marginal rugs refer to distributions of each individual axis regardless of whether a pairwise data point (commonly quantified peptide) is present.

**Supp. Figure S4.**
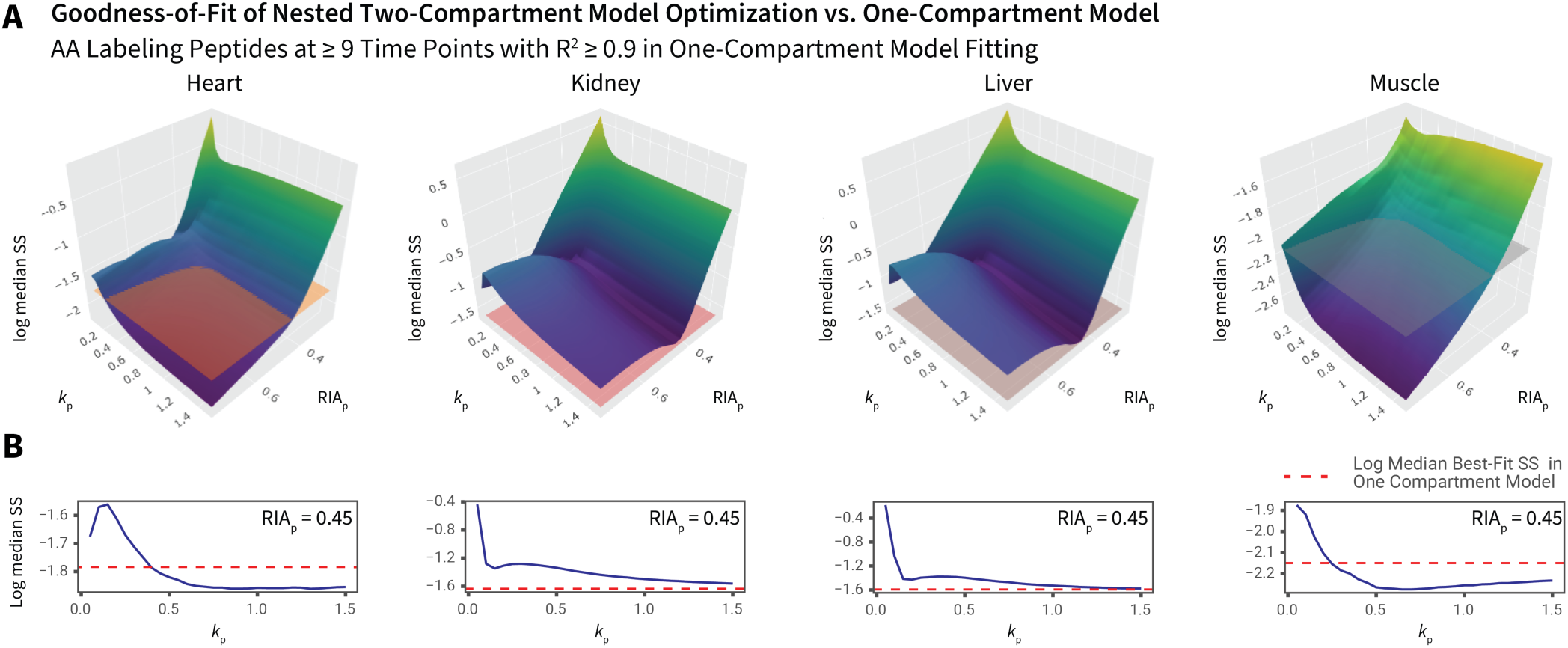
Determination of precursor RIA kinetics using proteome-wide nested optimization. **A.** We performed multiple rounds of two-compartment model curve-fitting by iterating through different *k*_p_ and plateau precursor RIA_p_ values from 0.05 to 2.0 at 0.05 increments (x-axis). Peptides quantified at ≥ 9 time points and fitted with R^2^ ≥ 0.9 in the one-compartment model were used. The median sums-of-squares of the residuals of fitting of each peptide time series in the two-compartment model in each tissue (z-axis) were compared to that from the one-compartment model (horizontal mesh). **B.** Corresponding two-dimensional cross-sections at various *k*_p_ values with asymptotic RIA_p_ fixed at 0.45. Red dash line: median sums-of-squares of peptide fitting in the simple exponential (one-compartment) model.

**Supp. Figure S5.**
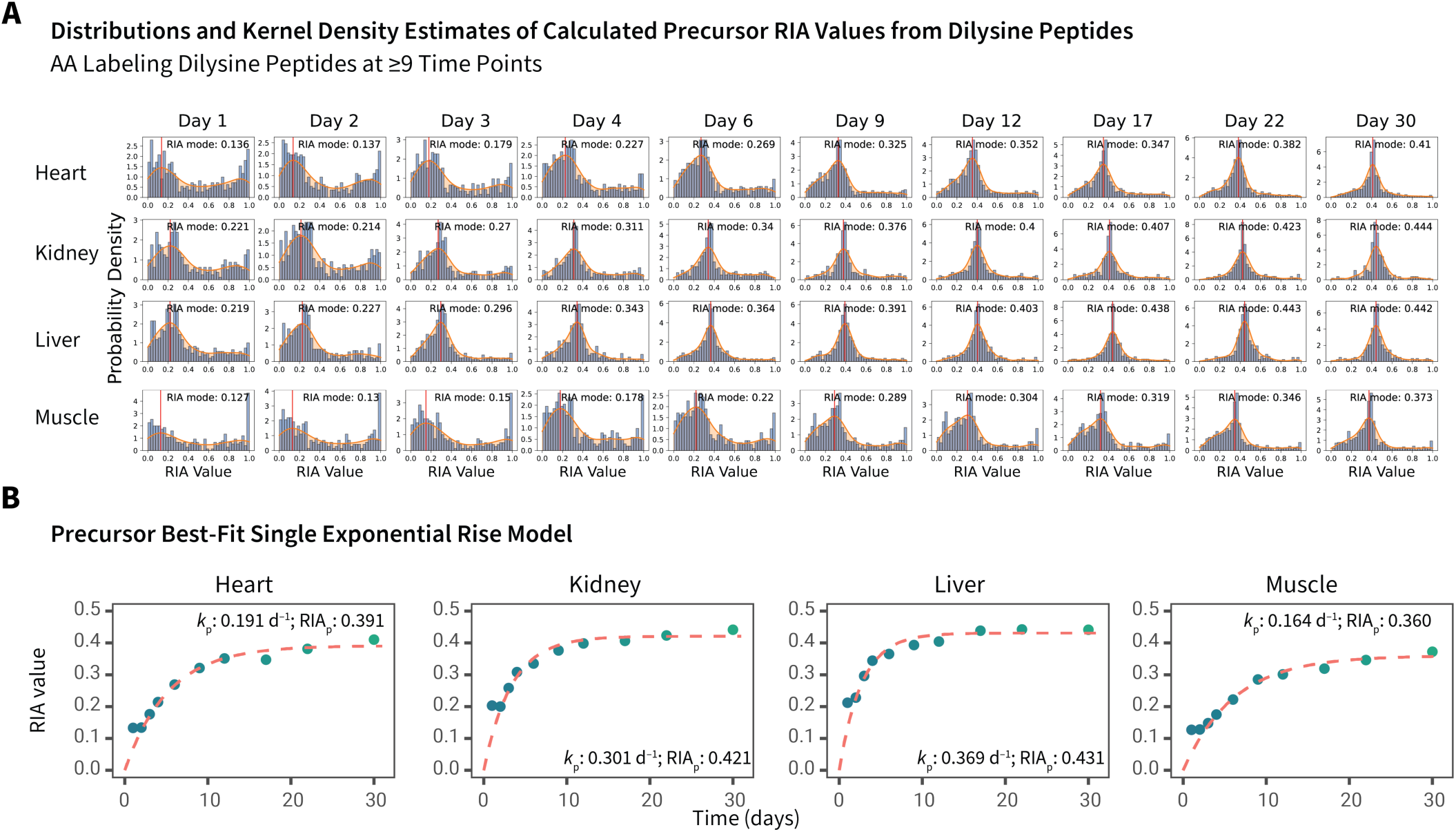
Determining AA precursor kinetics from dilysine peptides using Gaussian KDE. **A.** Distribution of calculated RIA values derived from mass isotopomer analysis of the intensities of the m_6_ and m_12_ peaks of peptides containing two lysine residues and quantified at ≥ 9 time points (bars). The best-estimate single tissue precursor RIA values for each time point for each tissue were derived from the modes of Gaussian kernel density estimations (red curve). **B.** The estimated tissue-specific precursor RIA values were fitted to a simple exponential model to derive the precursor rate constant *k*_p_ and asymptotic precursor RIA value RIA_p_.

**Supp. Figure S6.**
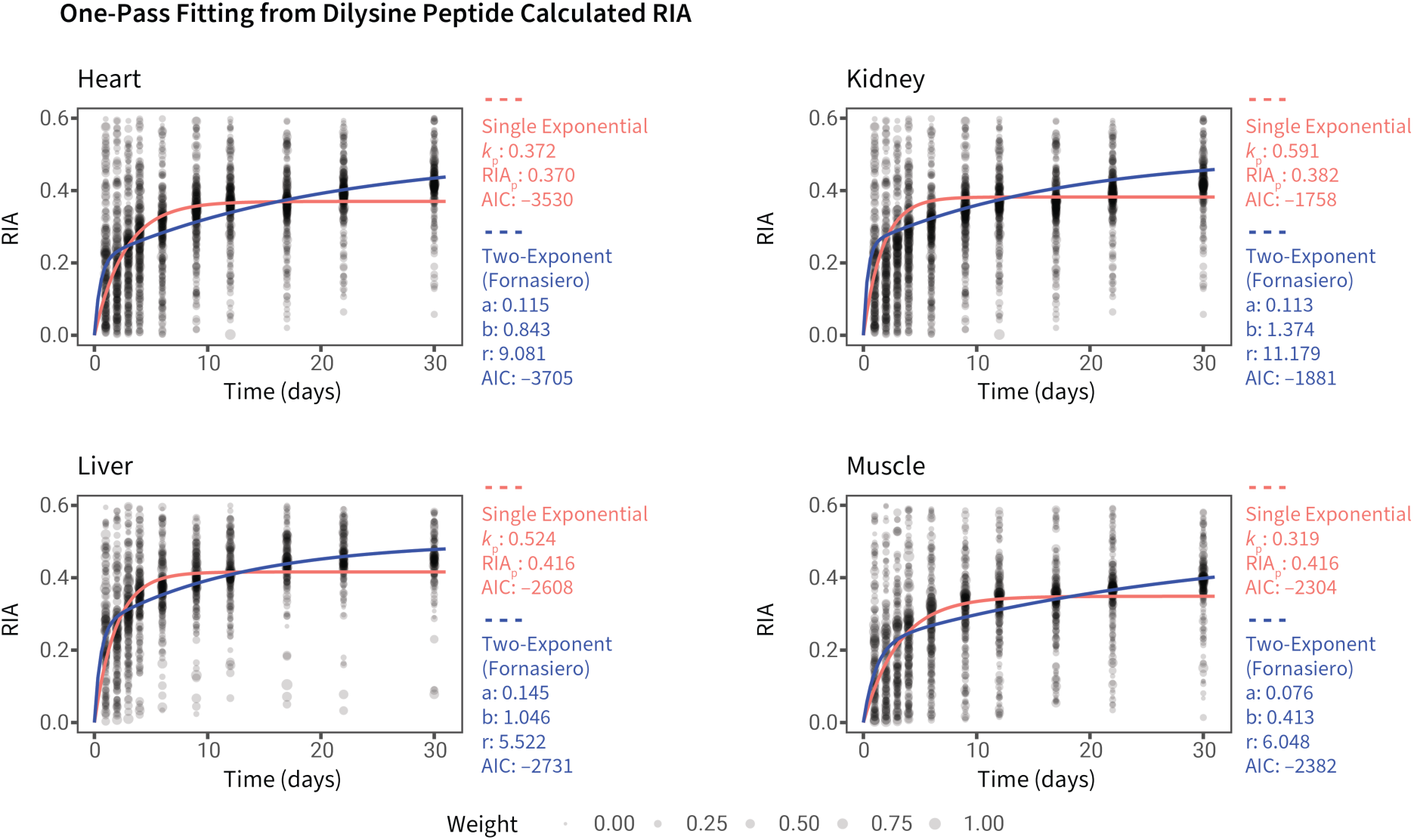
Determining AA precursor kinetics from dilysine peptides using weighted fitting. The distributions of calculated RIA values using the m_6_ and m_12_ peaks of dilysine peptides (y-axis) at each time point (x-axis) are shown across the four tissues, calculated as in Supplemental Figure S5. Data points with RIA between 0 and 0.6 are included and fitted directly to a simple exponential (red) model to find the best-fit *k*_p_ and plateau RIA_p_; or the double exponential (blue) model described in Fornasiero *et al.*^11^ to find the best-fit values for the parameters *a*, *b*, and *r* in the Fornasiero model. Weighted nonlinear least squares fitting was performed using the square of normalized peptide intensity of each data point as weight.

**Supp. Figure S7.**
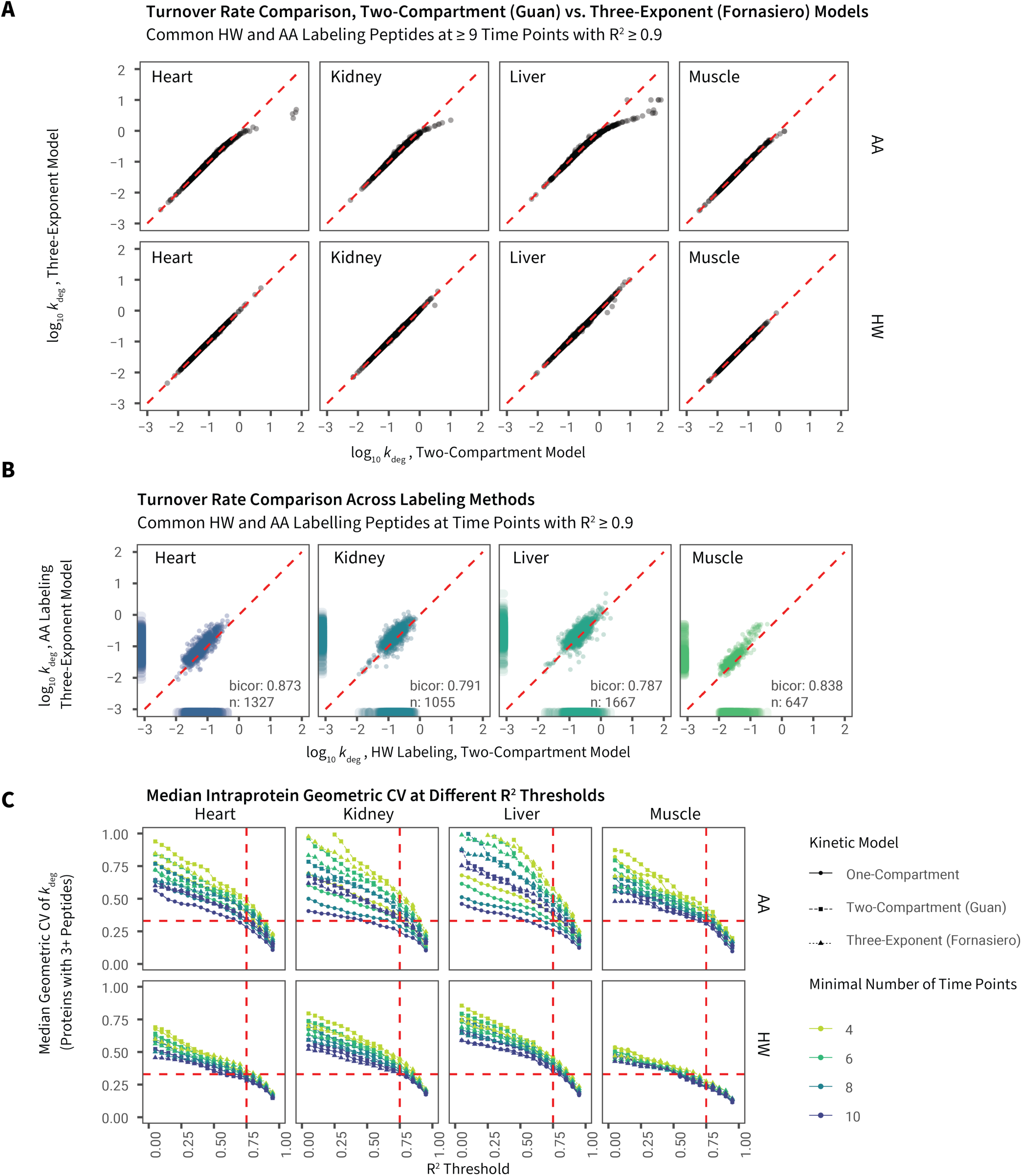
Comparison of the two-compartment and three-exponent models. **A.** Scatterplots showing the log_10_ peptide turnover rate constants in each tissue from HW and AA labeling derived using the Guan et al. two-compartment model (x-axis) and the Fornasiero three-exponent model (y-axis). Model parameters were derived using one-pass fitting to dilysine peptide RIA values as in Supplemental Figure S6. Each data point represents one common peptide quantified at ≥ 9 time points and fitted to each model as R^2^ ≥ 0.9. Red dashed lines: unity. **B.** Scatterplots comparing the log_10_ turnover rate constants in HW labeling (x-axis) derived using the two-compartment model with those in AA labeling (y-axis) derived using the three-exponent model. Numbers represent robust biweight midcorrelation (bicor) between HW and AA. **C.** Relationships between data variance as measured by intra-protein geometric coefficients of variation (CV) for proteins quantified with ≥ 3 peptides across multiple fitting R^2^ threshold (x-axis), with different time point multiplicity filters (color), and different kinetics models. Variance increased significantly when peptide time-series with lower fitting R^2^ were included, and the two compartment models showed higher variance than the simple exponential model.

**Supplemental Figure S8.**
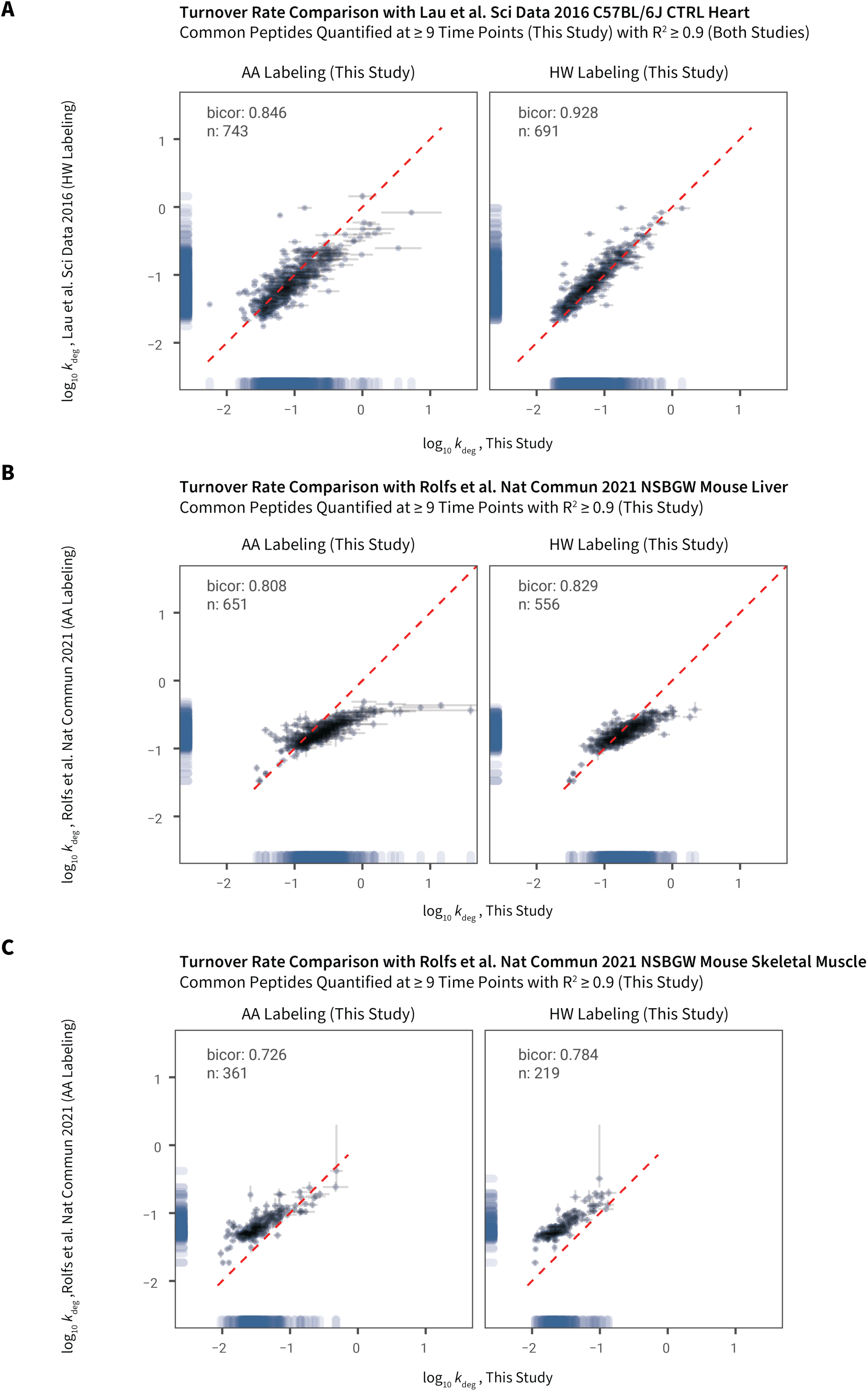
Comparison of turnover rate constants with a previous study. Scatterplots showing the log10 peptide turnover rates quantified with R^2^ ≥ 0.9 and at ≥ 9 time points in AA labeling (left) and HW labeling (right) in this study (x-axis) against the log10 peptide turnover rates in prior studies (y-axis): **A.** Peptides quantified with R^2^ ≥ 0.9 in C57BL/6J mouse heart in Lau et al. 2016 (HW labeling), vs. C57BL/6JOlaHsd mouse heart peptides in this study; **B.** reported peptides in NSBGW mouse liver in Rolfs et al. 2021, vs. C57BL/6JOlaHsd mouse liver peptides in this study; **C.** reported peptides in NSBGW skeletal (sternocleidomastoid) muscle in Rolfs et al. 2021, vs. C57BL/6JOlaHsd mouse skeletal muscle (pooled hindlimb) in this study. Bicor: biweight midcorrelation; n: number of compared peptides. Error bars: d*k*_deg_ of fitting. Dashed red line: unity.

## Supplemental Data

All Supplemental Data are available online on *figshare* at https://doi.org/10.6084/m9.figshare.17096636.v1

**Supplemental Data S1:** Table containing turnover rate constants of peptides

**Supplemental Data S2:** Fitted curves for common peptides in HW and AA labeling (≥ 9 time points; R^2^ ≥ 0.9) in the heart

**Supplemental Data S3:** Fitted curves for common peptides in HW and AA labeling (≥ 9 time points; R^2^ ≥ 0.9) in the kidney

**Supplemental Data S4:** Fitted curves for common peptides in HW and AA labeling (≥ 9 time points; R^2^ ≥ 0.9) in the liver

**Supplemental Data S5:** Fitted curves for common peptides in HW and AA labeling (≥ 9 time points; R^2^ ≥ 0.9) in the muscle

**Supplemental Data S6:** Table containing turnover rate constants of proteins

**Supplemental Data S7:** Protein-level fitted curves for common peptides in HW and AA labeling (≥ 9 time points; R^2^ ≥ 0.9) in the heart

**Supplemental Data S8:** Protein-level fitted curves for common peptides in HW and AA labeling (≥ 9 time points; R^2^ ≥ 0.9) in the kidney

**Supplemental Data S9:** Protein-level fitted curves for common peptides in HW and AA labeling (≥ 9 time points; R^2^ ≥ 0.9) in the liver

**Supplemental Data S10:** Protein-level fitted curves for common peptides in HW and AA labeling (≥ 9 time points; R^2^ ≥ 0.9) in the muscle

## References

(1) Doherty, M. K.; Hammond, D. E.; Clague, M. J.; Gaskell, S. J.; Beynon, R. J. Turnover of the Human Proteome: Determination of Protein Intracellular Stability by Dynamic SILAC. J. Proteome Res. 2009, 8 (1), 104–112. https://doi.org/10.1021/pr800641v.

(2) Pratt, J. M.; Petty, J.; Riba-Garcia, I.; Robertson, D. H. L.; Gaskell, S. J.; Oliver, S. G.; Beynon, R. J. Dynamics of Protein Turnover, a Missing Dimension in Proteomics. Mol. Cell. Proteomics MCP 2002, 1 (8), 579–591. https://doi.org/10.1074/mcp.m200046-mcp200.

(3) Zhang, T.; Price, J. C.; Nouri-Nigjeh, E.; Li, J.; Hellerstein, M. K.; Qu, J.; Ghaemmaghami, S. Kinetics of Precursor Labeling in Stable Isotope Labeling in Cell Cultures (SILAC) Experiments. Anal. Chem. 2014, 86 (22), 11334–11341. https://doi.org/10.1021/ac503067a.

(4) Hammond, D. E.; Claydon, A. J.; Simpson, D. M.; Edward, D.; Stockley, P.; Hurst, J. L.; Beynon, R. J. Proteome Dynamics: Tissue Variation in the Kinetics of Proteostasis in Intact Animals. Mol. Cell. Proteomics MCP 2016, 15 (4), 1204–1219. https://doi.org/10.1074/mcp.M115.053488.

(5) Claydon, A. J.; Thom, M. D.; Hurst, J. L.; Beynon, R. J. Protein Turnover: Measurement of Proteome Dynamics by Whole Animal Metabolic Labelling with Stable Isotope Labelled Amino Acids. Proteomics 2012, 12 (8), 1194–1206. https://doi.org/10.1002/pmic.201100556.

(6) Waterlow, J. C.; Garlick, P. J.; Millward, D. J. Protein Turnover in Mammalian Tissues and in the Whole Body; North-Holland Publishing Company, 1978.

(7) Dice, J. F.; Schimke, R. T. Turnover and Exchange of Ribosomal Proteins from Rat Liver. J. Biol. Chem. 1972, 247 (1), 98–111.

(8) Dice, J. F.; Dehlinger, P. J.; Schimke, R. T. Studies on the Correlation between Size and Relative Degradation Rate of Soluble Proteins. J. Biol. Chem. 1973, 248 (12), 4220–4228.

(9) Rahman, M.; Previs, S. F.; Kasumov, T.; Sadygov, R. G. Gaussian Process Modeling of Protein Turnover. J. Proteome Res. 2016, 15 (7), 2115–2122. https://doi.org/10.1021/acs.jproteome.5b00990.

(10) Nallasamy, S.; Palacios, H. H.; Setlem, R.; Colon Caraballo, M.; Li, K.; Cao, E.; Shankaran, M.; Hellerstein, M.; Mahendroo, M. Transcriptome and Proteome Dynamics of Cervical Remodeling in the Mouse during Pregnancy†. Biol. Reprod. 2021, 105 (5), 1257–1271. https://doi.org/10.1093/biolre/ioab144.

(11) Fornasiero, E. F.; Mandad, S.; Wildhagen, H.; Alevra, M.; Rammner, B.; Keihani, S.; Opazo, F.; Urban, I.; Ischebeck, T.; Sakib, M. S.; Fard, M. K.; Kirli, K.; Centeno, T. P.; Vidal, R. O.; Rahman, R.-U.; Benito, E.; Fischer, A.; Dennerlein, S.; Rehling, P.; Feussner, I.; Bonn, S.; Simons, M.; Urlaub, H.; Rizzoli, S. O. Precisely Measured Protein Lifetimes in the Mouse Brain Reveal Differences across Tissues and Subcellular Fractions. Nat. Commun. 2018, 9 (1), 4230. https://doi.org/10.1038/s41467-018-06519-0.

(12) Chepyala, S. R.; Liu, X.; Yang, K.; Wu, Z.; Breuer, A. M.; Cho, J.-H.; Li, Y.; Mancieri, A.; Jiao, Y.; Zhang, H.; Peng, J. JUMPt: Comprehensive Protein Turnover Modeling of In Vivo Pulse SILAC Data by Ordinary Differential Equations. Anal. Chem. 2021, 93 (40), 13495–13504. https://doi.org/10.1021/acs.analchem.1c02309.

(13) Bayram, H. L.; Claydon, A. J.; Brownridge, P. J.; Hurst, J. L.; Mileham, A.; Stockley, P.; Beynon, R. J.; Hammond, D. E. Cross-Species Proteomics in Analysis of Mammalian Sperm Proteins. J. Proteomics 2016, 135, 38–50. https://doi.org/10.1016/j.jprot.2015.12.027.

(14) Claydon, A. J.; Ramm, S. A.; Pennington, A.; Hurst, J. L.; Stockley, P.; Beynon, R. Heterogenous Turnover of Sperm and Seminal Vesicle Proteins in the Mouse Revealed by Dynamic Metabolic Labeling. Mol. Cell. Proteomics MCP 2012, 11 (6), M111.014993. https://doi.org/10.1074/mcp.M111.014993.

(15) Lau, E.; Cao, Q.; Ng, D. C. M.; Bleakley, B. J.; Dincer, T. U.; Bot, B. M.; Wang, D.; Liem, D. A.; Lam, M. P. Y.; Ge, J.; Ping, P. A Large Dataset of Protein Dynamics in the Mammalian Heart Proteome. Sci. Data 2016, 3, 160015. https://doi.org/10.1038/sdata.2016.15.

(16) Price, J. C.; Guan, S.; Burlingame, A.; Prusiner, S. B.; Ghaemmaghami, S. Analysis of Proteome Dynamics in the Mouse Brain. Proc. Natl. Acad. Sci. U. S. A. 2010, 107 (32), 14508–14513. https://doi.org/10.1073/pnas.1006551107.

(17) Lam, M. P. Y.; Wang, D.; Lau, E.; Liem, D. A.; Kim, A. K.; Ng, D. C. M.; Liang, X.; Bleakley, A. J.; Liu, C.; Tabaraki, J. D.; Cadeiras, M.; Wang, Y.; Deng, M. C.; Ping, P. Protein Kinetic Signatures of the Remodeling Heart Following Isoproterenol Stimulation. J. Clin. Invest. 2014, 124 (4), 1734–1744. https://doi.org/10.1172/JCI73787.

(18) Busch, R.; Kim, Y.-K.; Neese, R. A.; Schade-Serin, V.; Collins, M.; Awada, M.; Gardner, J. L.; Beysen, C.; Marino, M. E.; Misell, L. M.; Hellerstein, M. K. Measurement of Protein Turnover Rates by Heavy Water Labeling of Nonessential Amino Acids. Biochim. Biophys. Acta 2006, 1760 (5), 730–744. https://doi.org/10.1016/j.bbagen.2005.12.023.

(19) Previs, S. F.; Fatica, R.; Chandramouli, V.; Alexander, J. C.; Brunengraber, H.; Landau, B. R. Quantifying Rates of Protein Synthesis in Humans by Use of 2H2O: Application to Patients with End-Stage Renal Disease. Am. J. Physiol. Endocrinol. Metab. 2004, 286 (4), E665–672. https://doi.org/10.1152/ajpendo.00271.2003.

(20) Sadygov, R. G. Using Heavy Mass Isotopomers for Protein Turnover in Heavy Water Metabolic Labeling. J. Proteome Res. 2021, 20 (4), 2035–2041. https://doi.org/10.1021/acs.jproteome.0c00873.

(21) Sadygov, R. G.; Avva, J.; Rahman, M.; Lee, K.; Ilchenko, S.; Kasumov, T.; Borzou, A. D2ome, Software for in Vivo Protein Turnover Analysis Using Heavy Water Labeling and LC-MS, Reveals Alterations of Hepatic Proteome Dynamics in a Mouse Model of NAFLD. J. Proteome Res. 2018, 17 (11), 3740–3748. https://doi.org/10.1021/acs.jproteome.8b00417.

(22) Holmes, W. E.; Angel, T. E.; Li, K. W.; Hellerstein, M. K. Dynamic Proteomics: In Vivo Proteome-Wide Measurement of Protein Kinetics Using Metabolic Labeling. Methods Enzymol. 2015, 561, 219–276. https://doi.org/10.1016/bs.mie.2015.05.018.

(23) Guan, S.; Price, J. C.; Ghaemmaghami, S.; Prusiner, S. B.; Burlingame, A. L. Compartment Modeling for Mammalian Protein Turnover Studies by Stable Isotope Metabolic Labeling. Anal. Chem. 2012, 84 (9), 4014–4021. https://doi.org/10.1021/ac203330z.

(24) Rolfs, Z.; Frey, B. L.; Shi, X.; Kawai, Y.; Smith, L. M.; Welham, N. V. An Atlas of Protein Turnover Rates in Mouse Tissues. Nat. Commun. 2021, 12 (1), 6778. https://doi.org/10.1038/s41467-021-26842-3.

(25) Wiśniewski, J. R.; Zougman, A.; Nagaraj, N.; Mann, M. Universal Sample Preparation Method for Proteome Analysis. Nat. Methods 2009, 6 (5), 359–362. https://doi.org/10.1038/nmeth.1322.

(26) Broadhurst, D. I.; Kell, D. B. Statistical Strategies for Avoiding False Discoveries in Metabolomics and Related Experiments. Metabolomics 2006, 2 (4), 171–196. https://doi.org/10.1007/s11306-006-0037-z.

(27) Brown, M.; Dunn, W. B.; Ellis, D. I.; Goodacre, R.; Handl, J.; Knowles, J. D.; O’Hagan, S.; Spasić, I.; Kell, D. B. A Metabolome Pipeline: From Concept to Data to Knowledge. Metabolomics 2005, 1 (1), 39–51. https://doi.org/10.1007/s11306-005-1106-4.

(28) Mullard, G.; Allwood, J. W.; Weber, R.; Brown, M.; Begley, P.; Hollywood, K. A.; Jones, M.; Unwin, R. D.; Bishop, P. N.; Cooper, G. J. S.; Dunn, W. B. A New Strategy for MS/MS Data Acquisition Applying Multiple Data Dependent Experiments on Orbitrap Mass Spectrometers in Non-Targeted Metabolomic Applications. Metabolomics 2015, 11 (5), 1068–1080. https://doi.org/10.1007/s11306-014-0763-6.

(29) Wright Muelas, M.; Roberts, I.; Mughal, F.; O’Hagan, S.; Day, P. J.; Kell, D. B. An Untargeted Metabolomics Strategy to Measure Differences in Metabolite Uptake and Excretion by Mammalian Cell Lines. Metabolomics 2020, 16 (10), 107. https://doi.org/10.1007/s11306-020-01725-8.

(30) Hulstaert, N.; Shofstahl, J.; Sachsenberg, T.; Walzer, M.; Barsnes, H.; Martens, L.; Perez-Riverol, Y. ThermoRawFileParser: Modular, Scalable, and Cross-Platform RAW File Conversion. J. Proteome Res. 2020, 19 (1), 537–542. https://doi.org/10.1021/acs.jproteome.9b00328.

(31) UniProt Consortium. UniProt: The Universal Protein Knowledgebase in 2021. Nucleic Acids Res. 2021, 49 (D1), D480–D489. https://doi.org/10.1093/nar/gkaa1100.

(32) Eng, J. K.; Hoopmann, M. R.; Jahan, T. A.; Egertson, J. D.; Noble, W. S.; MacCoss, M. J. A Deeper Look into Comet--Implementation and Features. J. Am. Soc. Mass Spectrom. 2015, 26 (11), 1865–1874. https://doi.org/10.1007/s13361-015-1179-x.

(33) da Veiga Leprevost, F.; Haynes, S. E.; Avtonomov, D. M.; Chang, H.-Y.; Shanmugam, A. K.; Mellacheruvu, D.; Kong, A. T.; Nesvizhskii, A. I. Philosopher: A Versatile Toolkit for Shotgun Proteomics Data Analysis. Nat. Methods 2020, 17 (9), 869–870. https://doi.org/10.1038/s41592-020-0912-y.

(34) The, M.; MacCoss, M. J.; Noble, W. S.; Käll, L. Fast and Accurate Protein False Discovery Rates on Large-Scale Proteomics Data Sets with Percolator 3.0. J. Am. Soc. Mass Spectrom. 2016, 27 (11), 1719–1727. https://doi.org/10.1007/s13361-016-1460-7.

(35) Kösters, M.; Leufken, J.; Schulze, S.; Sugimoto, K.; Klein, J.; Zahedi, R. P.; Hippler, M.; Leidel, S. A.; Fufezan, C. PymzML v2.0: Introducing a Highly Compressed and Seekable Gzip Format. Bioinforma. Oxf. Engl. 2018, 34 (14), 2513–2514. https://doi.org/10.1093/bioinformatics/bty046.

(36) Virtanen, P.; Gommers, R.; Oliphant, T. E.; Haberland, M.; Reddy, T.; Cournapeau, D.; Burovski, E.; Peterson, P.; Weckesser, W.; Bright, J.; van der Walt, S. J.; Brett, M.; Wilson, J.; Millman, K. J.; Mayorov, N.; Nelson, A. R. J.; Jones, E.; Kern, R.; Larson, E.; Carey, C. J.; Polat, İ.; Feng, Y.; Moore, E. W.; VanderPlas, J.; Laxalde, D.; Perktold, J.; Cimrman, R.; Henriksen, I.; Quintero, E. A.; Harris, C. R.; Archibald, A. M.; Ribeiro, A. H.; Pedregosa, F.; van Mulbregt, P.; SciPy 1.0 Contributors. SciPy 1.0: Fundamental Algorithms for Scientific Computing in Python. Nat. Methods 2020, 17 (3), 261–272. https://doi.org/10.1038/s41592-019-0686-2.

(37) Berglund, M.; Wieser, M. E. Isotopic Compositions of the Elements 2009 (IUPAC Technical Report). Pure Appl. Chem. 2011, 83 (2), 397–410. https://doi.org/10.1351/PAC-REP-10-06-02.

(38) Commerford, S. L.; Carsten, A. L.; Cronkite, E. P. The Distribution of Tritium among the Amino Acids of Proteins Obtained from Mice Exposed to Tritiated Water. Radiat. Res. 1983, 94 (1), 151–155.

(39) Langfelder, P.; Horvath, S. WGCNA: An R Package for Weighted Correlation Network Analysis. BMC Bioinformatics 2008, 9, 559. https://doi.org/10.1186/1471-2105-9-559.

(40) Pedersen, T. L. Ggplot2, 2021.

(41) Maag, J. L. V. Gganatogram: An R Package for Modular Visualisation of Anatograms and Tissues Based on Ggplot2. F1000Research 2018, 7, 1576. https://doi.org/10.12688/f1000research.16409.2.

(42) Kassambara, A. Ggpubr: “ggplot2” Based Publication Ready Plots, 2021.

(43) Sievert, C. Plotly, 2021.

(44) Cavaggioni, A.; Mucignat-Caretta, C. Major Urinary Proteins, Alpha(2U)-Globulins and Aphrodisin. Biochim. Biophys. Acta 2000, 1482 (1–2), 218–228. https://doi.org/10.1016/s0167-4838(00)00149-7.

(45) Hurst, J. L.; Beynon, R. J.; Armstrong, S. D.; Davidson, A. J.; Roberts, S. A.; Gómez-Baena, G.; Smadja, C. M.; Ganem, G. Molecular Heterogeneity in Major Urinary Proteins of Mus Musculus Subspecies: Potential Candidates Involved in Speciation. Sci. Rep. 2017, 7, 44992. https://doi.org/10.1038/srep44992.

(46) Roberts, S. A.; Prescott, M. C.; Davidson, A. J.; McLean, L.; Beynon, R. J.; Hurst, J. L. Individual Odour Signatures That Mice Learn Are Shaped by Involatile Major Urinary Proteins (MUPs). BMC Biol. 2018, 16 (1), 48. https://doi.org/10.1186/s12915-018-0512-9.

(47) Hammond, D. E.; Kumar, J. D.; Raymond, L.; Simpson, D. M.; Beynon, R. J.; Dockray, G. J.; Varro, A. Stable Isotope Dynamic Labeling of Secretomes (SIDLS) Identifies Authentic Secretory Proteins Released by Cancer and Stromal Cells. Mol. Cell. Proteomics MCP 2018, 17 (9), 1837–1849. https://doi.org/10.1074/mcp.TIR117.000516.

(48) Hubrecht, R. C.; Carter, E. The 3Rs and Humane Experimental Technique: Implementing Change. Anim. Open Access J. MDPI 2019, 9 (10), E754. https://doi.org/10.3390/ani9100754.

